# MicroRNA-132 provides neuroprotection for tauopathies via multiple signaling pathways

**DOI:** 10.1101/258509

**Authors:** Rachid El Fatimy, Shaomin Li, Zhicheng Chen, Tasnim Mushannen, Sree Gongala, Zhiyun Wei, Darrick T. Balu, Rosalia Rabinovsky, Adam Cantlon, Abdallah Elkhal, Dennis J. Selkoe, Kai C. Sonntag, Dominic M. Walsh, Anna M. Krichevsky

## Abstract

MicroRNAs (miRNA) regulate fundamental biological processes, including neuronal plasticity, stress response, and survival. Here we describe a neuroprotective function of miR-132, the miRNA most significantly down-regulated in Alzheimer’s disease. miR-132 protects mouse and human wild-type neurons and more vulnerable Tau-mutant primary neurons against amyloid β-peptide (Aβ) and glutamate excitotoxicity. It lowers the levels of total, phosphorylated, acetylated, and cleaved forms of Tau implicated in tauopathies, promotes neurite elongation and branching, and reduces neuronal death. Similarly, miR-132 attenuates PHF Tau pathology and neurodegeneration and enhances long-term potentiation in the P301S Tau transgenic mice. The neuroprotective effects are mediated by direct regulation of the Tau modifiers acetyltransferase EP300, kinase GSK3β, RNA-binding protein Rbfox1, and proteases Calpain 2 and Caspases 3/7. These data suggest miR-132 as a master regulator of neuronal health and indicate that miR-132 supplementation could be of therapeutic benefit for the treatment of Tau-associated neurodegenerative disorders.

## Introduction

Alzheimer’s disease (AD) is a progressive neurodegenerative disorder typified by profound synaptic loss, brain atrophy and the presence of extracellular plaques composed of amyloid β-protein (Aβ), and intracellular neurofibrillary tangles (NFTs) formed by hyperphosphorylated Tau^1,2^. NFTs are also pathogenomic for a range of disorders in which Tau deposits occur in the absence of plaques. The major primary tauopathies include: frontotemporal dementia (FTD), progressive supranuclear palsy (PSP), Pick’s disease, and corticobasal degeneration. New therapeutic strategies and targets are desperately needed to treat these devastating diseases^3^.

miRNAs are small regulatory molecules that post-transcriptionally repress gene expression and thereby regulate diverse biological processes, including neuronal differentiation, plasticity, survival, and regeneration^4,5^. miRNAs are often considered as determinants of cell fate and are also increasingly acknowledged as prime regulators involved in various brain pathologies ranging from neurodevelopmental disorders to brain tumors, to neurodegenerative diseases^6,7^. Given its immense complexity, the brain expresses more distinct miRNA species than any other organ in the body^8^, with specific miRNAs being highly enriched in certain cell types of the brain, e.g. developing or mature cortical neurons. Early studies reported that deficiency of Dicer, the key ribonuclease in miRNA biogenesis, resulted in progressive miRNA loss, death of Purkinje neurons and cerebellar degeneration^9,10,11^. Several neuronal miRNAs have been directly linked to the regulation of key factors involved in AD, including APP and Aβ. production and clearance^12^. Although multiple lines of evidence suggest ^that^ miRNAs may contribute to the progression of neurodegenerative diseases, the complexity of miRNA regulation in targeting many genes and pathways simultaneously raised concerns about their therapeutic utility as targetable molecules.

One of the most abundant brain-enriched miRNAs is miR-132, which plays a key role in both neuron morphogenesis and plasticity. miR-132, transcribed by the activity-dependent transcription factor CREB, modulates dendritic plasticity, growth and spine formation in response to a variety of signaling pathways^13,14^. Deletion of the miR-132 locus decreases dendritic arborization, length and spine density, impairs integration of newborn neurons, and reduces synapse formation in the adult hippocampus^15,16^. miR-132 also regulates axon extension through the Ras GTPase activator Rasa1^17^. Similarly, miR-132 inhibition induces apoptosis in cultured cortical and hippocampal primary neurons via PTEN/AKT/FOXO3 signaling^18^. Notably, miR-132-deficient mice exhibit Tau hyperphosphorylation, aggregation, and decreased memory – all of which are hallmarks of AD^19,20^. Deletion of miR-132 also fosters Aβ production and plaque accumulation in a triple transgenic AD model^21^. Importantly, multiple studies have shown that miR-132 is the most downregulated miRNA in postmortem AD brain with reductions in miR-132 occurring before neuronal loss, and associated with progression of both amyloid and Tau pathology^18,19,21,22,23,24^.

We hypothesized that supplementation of miR-132 activity may protect against AD and other tauopathies. In support of this idea, we report results of a high-content miRNA screen performed on primary mouse and human neurons treated with either an AD specific insult (Aβ) or excitotoxic levels of glutamate. Among the miRNAs expressed in brain miR-132 exhibits the strongest neuroprotective activity against both Aβ and glutamate. Furthermore, overexpression of miR-132 reduced phosphorylated, acetylated, and cleaved forms of Tau in primary neurons, as well as Tau pathology and caspase-3-dependent apoptosis in PS19 (Tau^P301S^) mice. Functionally, miR-132 overexpression enhanced long-term potentiation (LTP) in WT mice and rescued the impairment of LTP seen in PS19 mice. These results suggest that miR-132 replacement could provide neuroprotection and therapeutic value for Tau-associated neurodegenerative disorders, including AD and FTD.

## RESULTS

### A miRNA screen identifies miR-132 as strongly neuroprotective against Aβ and glutamate excitotoxicity in primary neurons

To identify endogenous miRNAs with neuroprotective properties, we performed a screen on 63 conserved neuronal miRNAs that together account for more than 90% of all miRNA expressed in the adult mouse brain^25^. Individual miRNAs were inhibited in mouse primary hippocampal and cortical neurons by specific locked nucleic acid (LNA)-based antisense oligonucleotide *inhibitors* (anti-miRNAs). Alternatively, cells were transfected with non-targeting control oligonucleotides of the same chemistry. The neurons were either transfected at DIV7 and exposed to glutamate (100 μM) two days later, or transfected at DIV19 followed by the treatment with toxic Aβ (15 μM) (Figure S1). Based on the general consensus that aggregation of Aβ is required for toxicity, we employed a partially aggregated preparation of Aβ (1–42) that contained both amyloid fibrils and Aβ monomer and is referred to as ½tmaxAβ^26,27^. Metabolic activity and cell viability were assessed by WST-1 assays three days after glutamate or Aβ exposure, and the effects of miRNA inhibitors normalized to mock and control oligonucleotide (Fig. 1A). We observed that inhibition of specific miRNAs protected against glutamate (e.g., anti-let-7g,i), Aβ toxicity (e.g., anti-miRNA-7), or both toxic stimuli (e.g., let-7c,e) (Fig. 1A). In contrast, inhibition of other miRNAs (e.g., miR-107, miR-30a, miR-29, miR-212-3p, and miR-132-3p) exacerbated Aβ and glutamate toxicities (Fig. 1A), suggesting neuroprotective functions for this group of miRNAs.

**Figure 1.**
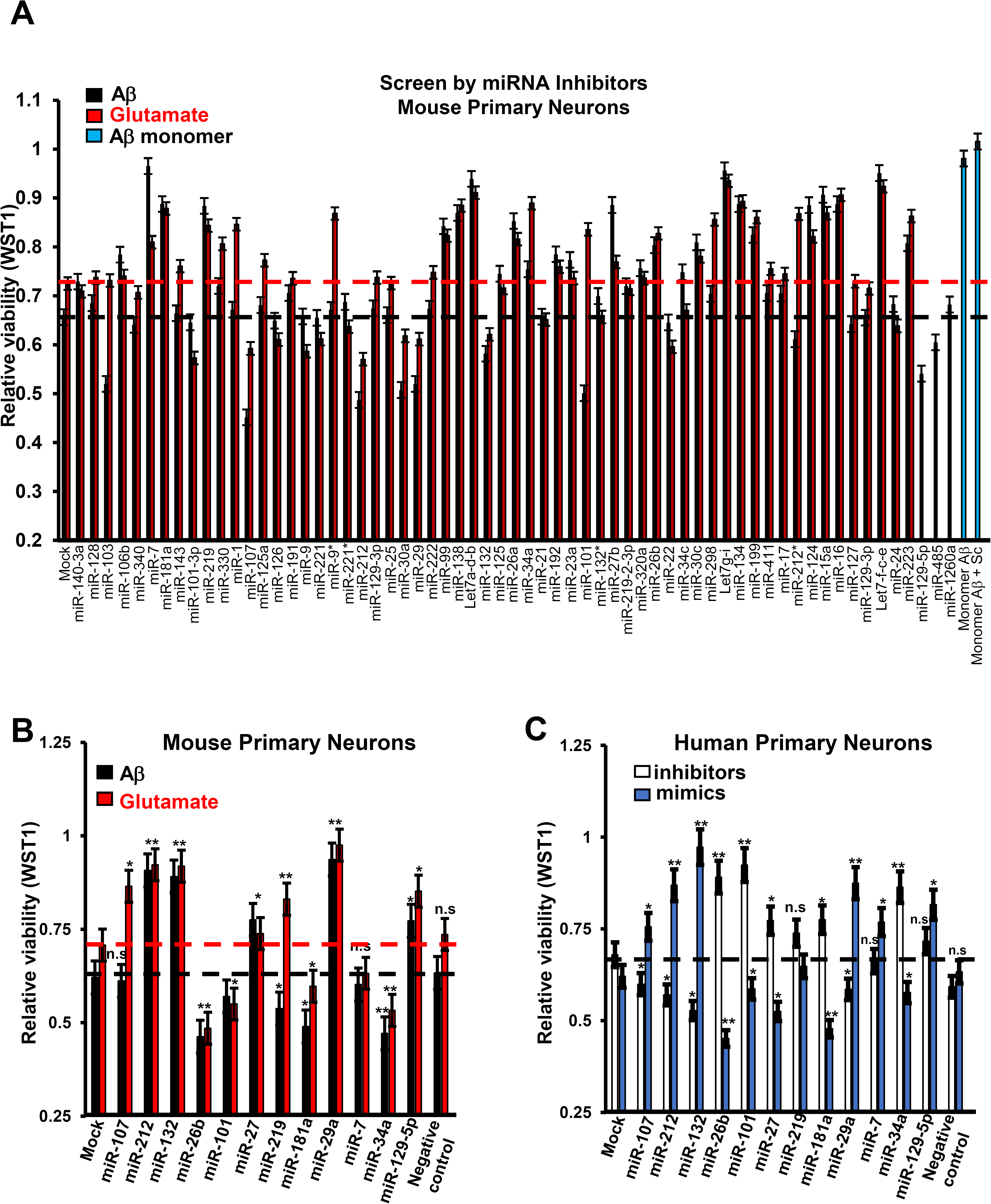
A screen in primary neurons identifies miRNAs protecting or exacerbating Aβ and glutamate toxicity. Primary neurons were transfected with either miRNA oligonucleotide inhibitors (anti-miRNA) or mimics (miRNA), and treated with Aβ (15 μM), glutamate (100 μM) or vehicle. WST1 assays were performed three days later, and results normalized to transfected, vehicle-treated neurons (see also Figure S1). Six wells per condition were analyzed. (**A**) Relative viability of mouse hippocampal neurons transfected with the indicated sequence-specific anti-miRNAs. The value of “1” corresponds to the viability of untreated controls neurons. Treatment with non-toxic Aβ monomer does not reduce the viability (shown by two bars on the right side of the panel). The horizontal black and red lines mark the viability of untransfected neurons treated with Aβ or glutamate, respectively. (**B**) Relative viability of mouse hippocampal neurons transfected with the indicated miRNA *mimics*. (**C**) Relative viability of *human* primary cortical neurons pre-transfected with either miRNA inhibitors or mimics and then treated with Aβ. miR-132 mimic induced the most significant increase of neuronal survival in both mouse and human cultures (*P <0.05 and **P <0.01), n= 6, Student’s t-test, control versus miR-132 inhibitor or mimic condition). Graphical data are shown as mean +/- SEM.

To validate miRNAs modulating neuronal survival in these conditions, we transfected mouse hippocampal neurons with oligonucleotide *mimics* of the top 12 hits and then added Aβ or glutamate. As expected, most miRNAs whose inhibitors decreased the viability in the primary screen, appeared neuroprotective when their mimics were applied (miR-132-3p, miR-212-3p, miR-129-5p, and miR-29a-5p in Fig. 1B). Conversely, the miRNAs whose inhibitors were neuroprotective, exacerbated the toxicity (miR-26b, -34a in Fig. 1B). Similar experiments with both miRNA inhibitors and mimics were also performed on *human* primary cortical neurons stressed with Aβ (Fig. 1C). The results in the rodent and human cultures were in good agreement for the 12 miRNAs tested, and in both cases miR-132(-3p) was identified as the most neuroprotective and miR-26b as the most “neurotoxic” miRNA (Fig. 1B, C). Notably, miR-132 mimic increased the survival of both human and mouse neurons treated with Aβ by ~20% (Fig. 1A, B, C). We also observed that Aβ-induced Tau hyperphosphorylation, which is essential for Aβ toxicity in neurons^28,29^, was restrained by miR-132 overexpression (Fig. 2A, B).

**Figure 2.**
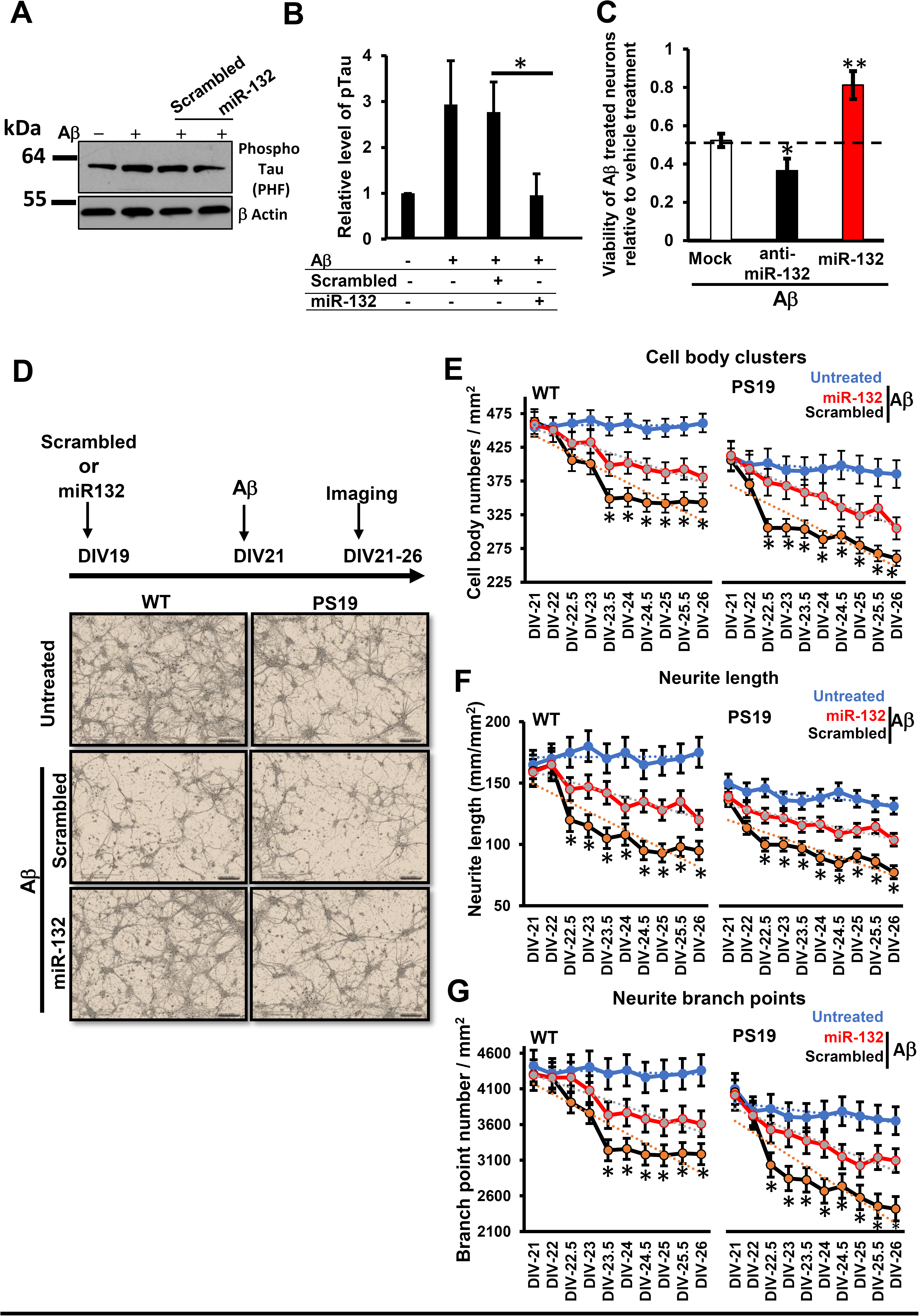
MiR-132 recues morphology and improves health of WT and Tau P301S neurons treated with toxic Aβ species. **(A)** Western blot analysis of phospho-Tau (PHF1) in mouse primary neurons transfected with miR-132 mimic and treated with Aβ demonstrates that miR-132 reduces PHF1 levels. (**B**) Quantification of three Western blot experiments similar to that shown in (A). (**C**) Relative viability of primary PS19/P301S neurons transfected with either anti-miR-132, or miR-132 mimic oligonucleotides, and treated with Aβ. The data were normalized to the viability of untreated cells. (**D**) Live-cell imaging of WT and PS19 neurons, and representative images shown for DIV26. Quantification of the (**E**) cell body clusters, (**F**) neurite length, and (**G**) number of neurite branch points, for 18 images per condition taken between days 23-26, derived from 8 primary cultures. Graphical data are shown as mean +/- SEM, n = 18 per condition, *P <0.005, Student’s t-test.

### Overexpression of miR-132 preserves cell body clusters and neurite integrity in WT and PS19 neurons treated with ½ t_max_ Aβ

To further investigate the protective effects of miR-132 overexpression *in vitro* and *in vivo* we used a PS19 tau transgenic mouse line which expresses human 1N4R Tau bearing the P301S mutation associated with FTD^30^. PS19 primary neurons were transfected with either anti-miR-132, miR-132 mimic, or control oligonucleotides, and then treated with aggregated Aβ. As with WT neurons, in PS19 primary neurons, the miR-132 mimic protected against Aβ while the anti-miR-132 exacerbated sensitivity to Aβ (Fig. 2C). To investigate the effects of miR-132 overexpression on neuronal morphology in normal versus Aβ-stressed cultures we imaged live cells over a 5-day interval (Fig. 2D). Of note, naïve PS19 cultures appeared less healthy than WT cultures, exhibited fewer and more clustered cell bodies, and had shorter and less branched neurites. In both WT and PS19 neurons stressed with ½t_max_ Aβ, transfection of miR-132 mimic increased the number of healthy cell bodies, neurite length and branch points versus neurons transfected with scrambled oligonucleotides (Fig. 2E, F, G). These significant effects demonstrate that under stress or toxic conditions miR-132 rescues neuritic loss and helps maintain neuronal integrity in both WT and mutant Tau neurons (Fig. 2E, F, G).

### miR-132 reduces the levels of total and post-translationally modified forms of Tau, its cleavage and release in PS19 neurons

miR-132 downregulation is significantly associated with human Tau pathology^22,24^, and its genetic deficiency increases Tau expression, phosphorylation, and aggregation in 3×Tg AD transgenic mice^20^. We therefore investigated the effects of miR-132 mimic on Tau metabolism which in disease is dysregulated at multiple levels. Tau hyperphosphorylation is a hallmark of most tauopathies^2^, but several other post-translational modifications are also common. For instance, acetylation at Lys174 (K174) which hinders the interaction between Tau and microtubules and is thought to foster Tau accumulation and aggregation^31,32,33^. Also, Tau can undergo proteolytic cleavages which generate fragments^34,35,36^(Fig. 3A) some of which are prone to aggregation and are suggested to be toxic^37,38^.

**Figure 3.**
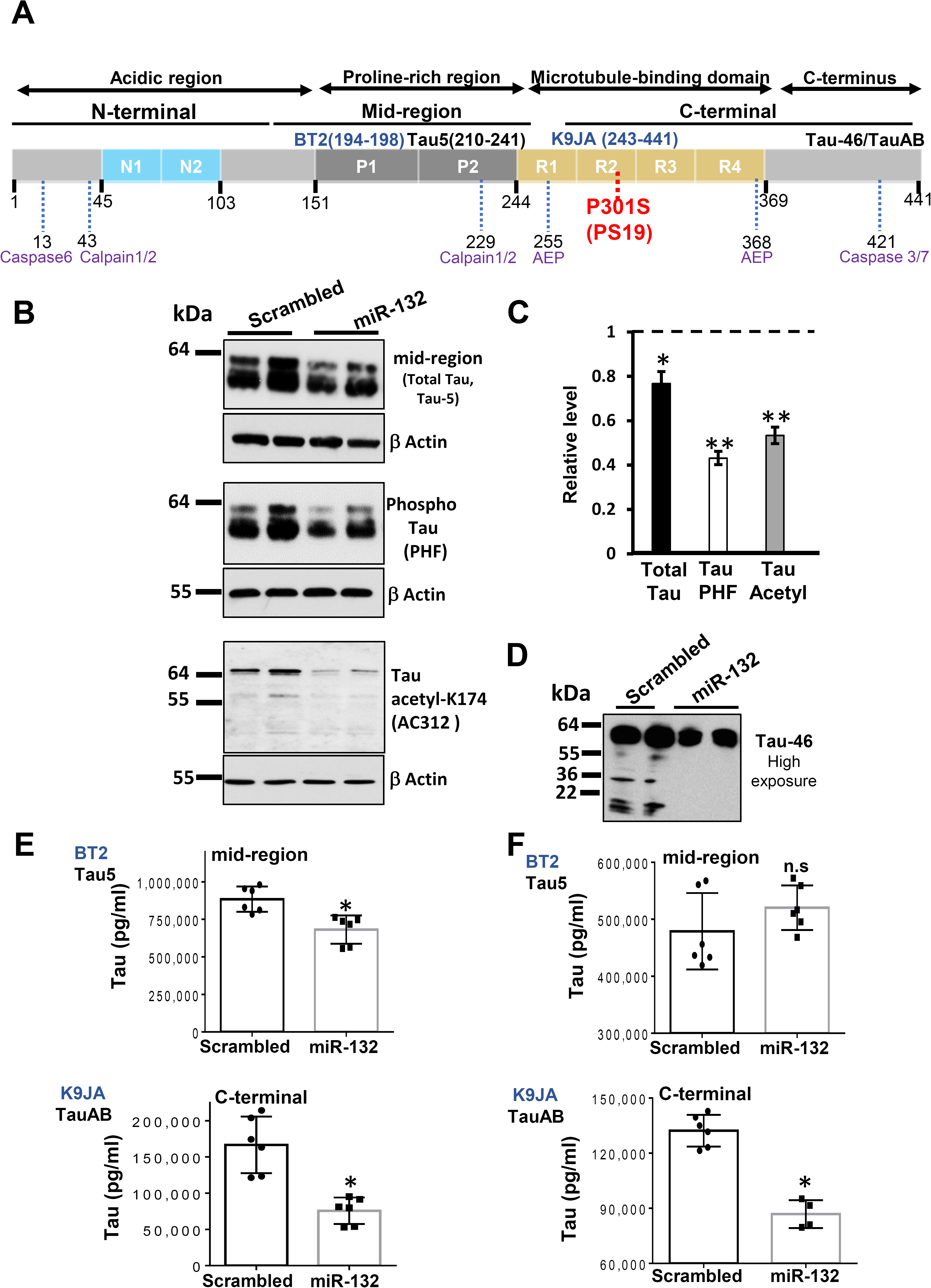
MiR-132 reduces the levels of total Tau, its phosphorylated and acetylated forms, Tau cleavage and release of a C-terminal fragment. **(A)** A schematic presentation of Tau protein domains, positions of the epitopes recognized by the antibodies utilized in this study, and proteolytic sites of major proteases. The longest human CNS isoform, Tau 441, is depicted. (**B-E**) Primary PS19 cortical neurons were transfected with either miR-132 mimics or scramble control oligonucleotides, and protein lysates collected 48-hours post-transfection and analyzed. β-actin served as a loading control for all Western blots. (**B**) Western blots analyses demonstrate the effects of miR-132 mimic on the levels of total Tau, and its phosphorylated and acetylated forms. Primary antibodies used are indicated in the parentheses. (**C**) Quantification of the levels of total, phosphorylated, and acetylated Tau in neurons transfected with miR-132, plotted relative to the levels in neurons transfected with scrambled oligonucleotides. *P < 0.05, and **P <0.005, n = 12 cultures per condition, t-test. (**D**) Western blots analysis demonstrates that miR-132 reduces Tau fragmentation. (**E**) Quantification of mid-region (left panel) and C-terminal (right panel) containing forms of Tau in neuronal lysates, measured by the BT2-Tau5 and K9JA-TauAB ELISAs (n=6 for each group). (**F**) Quantification of mid-region (left panel) and C-terminal (right panel) containing forms of Tau in neuronal conditioned medium, measured by ELISA assays (n=6 for each group). \**P* < 0.005, Student’s t-test, and *n* = 3. n.s, not significant. All graphical data are shown as mean +/- SEM. See also Figure S2.

Western blot analysis revealed that PS19 primary neurons transfected with miR-132 mimic exhibited slightly reduced levels of total Tau (Fig. 3B) and pronounced reduction of Tau phosphorylated at Ser396 and Ser404 (PHF1 epitope), and acetylated at K174 (Fig. 3B)^35^. Comparable results were obtained for Tau acetylated at the epitope K274 (Figure S2). Quantification of three independent experiments indicated that the reduction of post-translationally modified Tau isoforms was more pronounced than that of total Tau (Fig. 3C). These data indicate that the observed decrease in the levels of phosphorylated and acetylated Tau was not merely a consequence of reduced total Tau. Additional analysis of major Tau fragments in PS19 neurons using an antibody against the C-terminal region (Tau46) revealed that miR-132 mimic also reduced the levels of ~36kDa and ~17 kDa Tau fragments (Fig. 3D), the latter being previously characterized as a potentially neurotoxic fragment(s) produced by Calpain 2 and Caspase 3 proteolytic activities, significant amounts of which were found in the brains of patients with tauopathies^27,35,39,40,41^. Two distinct sandwich ELISA assays, one based on Tau mid-region detection, reflective of total Tau, and the other based on the detection of C-terminal fragments (capture and detection antibodies are illustrated in Fig. 3A) confirmed that miR-132 produces a small (~20%) but significant reduction of the levels of mid-region-containing Tau, and stronger reduction of the levels of C-terminal-containing Tau (Fig. 3E).

It is now widely appreciated that Tau exists both inside and outside of neurons^42,43,44^. Under normal circumstances the majority of extracellular Tau is C-terminal truncated^44,45,46,47^, but it has been speculated that in disease MTBR-containing forms of tau are released and may contribute to the seeding and spreading of tau aggregates^48^. Since miR-132 diminishes C-terminal Tau fragments inside primary neurons (Fig. 3D), we next asked whether it may also affect the release of Tau. Notably, although the concentrations of extracellular mid-region–containing tau were unaffected by miR-132 (Fig. 3F, left), the levels of extracellular C-terminal-containing secreted fragments were strongly reduced (Fig. 3F, right). Collectively, these results indicate that Tau homeostasis is regulated by miR-132 at several levels, including the regulation of its post-translational modifications, cleavage, and release from neurons.

### miR-132 directly targets the Tau modifiers Rbfox1, GSK3β, EP300, and Calpain 2

To investigate miR-132 regulation of Tau metabolism and discover among putative miR-132 targets those that modify Tau, we systematically applied several target prediction algorithms. Tau mRNA itself has been proposed as a direct target of miR-132^20^; however, previous work failed to confirm this in human neural cells^36^. Here, we identified several molecules that directly link miR-132 to Tau. mRNA for Glycogen Synthase Kinase-3 β (GSK3β) that contributes to pathologic Tau hyper-phosphorylation in AD^49,50,51,52^, contains two putative miR-132 binding sites within its 3’UTR. Transfections of primary neurons with miR-132 mimic reduced GSK3β amounts at both the mRNA and protein levels (Fig. 4A, B). Furthermore, a luciferase reporter containing the wild-type GSK3β 3′ UTR was repressed by the miR-132 mimic and up-regulated by the miR-132 inhibitor. These effects on the reporter were abrogated by a mutation of the predicted conserved binding sites at the position 2644-2650 (Fig. 4C, D). These data provide evidence that miR-132 directly binds to the GSK3β mRNA and reduces its expression, thereby lowering the levels of GSK3β-mediated phospho-Tau.

**Figure. 4.**
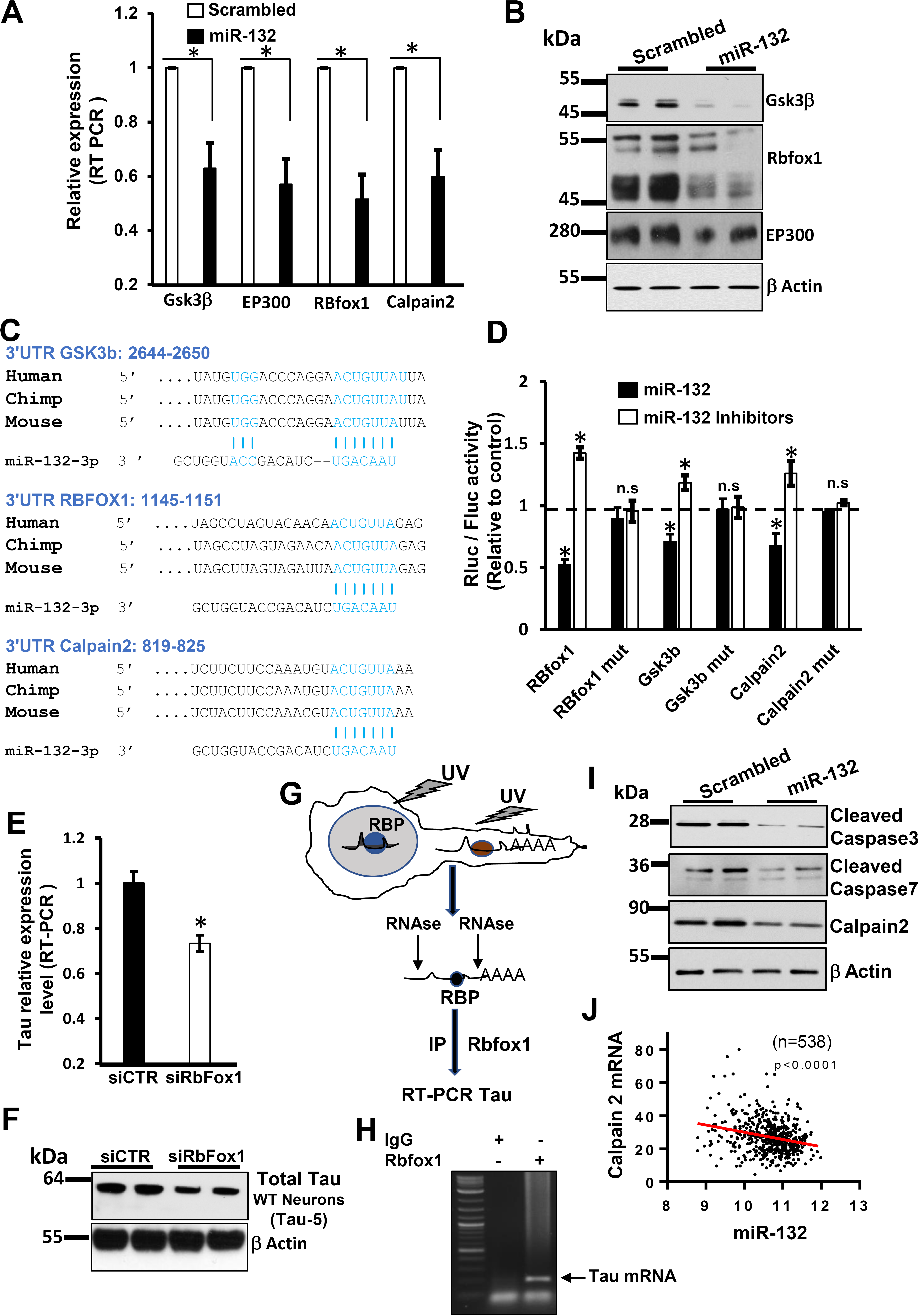
MiR-132 directly targets the Tau modifiers Rbfox1, GSK3β, EP300, and Calpain 2. **(A)** qRT-PCR analysis demonstrates the effects of miR-132 mimic on the expression levels of GSK3β, EP300, Rbfox1 and Calpain 2 mRNAs in WT neurons (*P <0.005, n=6, Student’s t-test). (**B**) The effects of miR-132 mimic on protein levels of GSK3β, Rbfox1 and EP300. (**C**) Predicted miR-132 binding sites within the *GSK3β, Calpain 2, and RBFOX1 3′* UTRs. (**D**) Luciferase reporter assays demonstrate that *RBFOX1, Gsk3β* and *Calpain 2* are direct miR-132 targets, modulated inversely by the miR-132 mimic and inhibitor. Relative activity of luciferase constructs bearing either wild-type (wt) or mutant (mut) *RBFOX1, GSK3β, or Calpain 2 3′* UTRs in neural cells co-transfected with miR-132 mimic or inhibitor is presented as renilla/firefly luminescence units (RLU). The data were normalized to the effects of the corresponding control oligonucleotides (set as “1”). \**P* < 0.005 and *n* = 3; n.s., not significant. Graphical data are shown as mean +/- SEM. (**E**) qRT-PCR and (**F**) Western Blot analyses demonstrate that RbFox1 silencing reduces the levels of total Tau mRNA and protein, respectively, in WT neurons. *P <0.01, n=3, student t-test. (**G**) Schematic presentation of the iCLIP experimental workflow. (**H**) iCLIP demonstrates that total Tau mRNA is co-immunoprecipitated with Rbfox1 protein from WT neurons. The immunoprecipitated RNA was amplified by RT-PCR and visualized by gel electrophoresis. (**I**) Western blot analyses demonstrate the effects of miR-132 mimic on the levels of cleaved caspase-3, caspase-7 and calpain 2. (**J**) Expression correlation between Calpain 2 mRNA and miR-132 in human prefrontal cortex, from the two cohorts of aging brains (n=538, p<0.0001, 61% meet criteria for pathologic AD by NIA Reagan criteria). The mRNA and miRNA datasets were downloaded from https://www.synapse.org/#!Synapse:syn3219045. See also Figure S3.

Acetyltransferase EP300 is the major acetylase of Tau at K174 implicated in Tau aggregation and neurodegeneration in AD^32^, and has previously been reported as one of the miR-132 targets contributing to its pro-survival/anti-apoptotic function^18^. Indeed, miR-132 mimics reduced expression of EP300, at both mRNA and protein levels in primary neurons (Fig. 4A, B). These data indicate that the observed miR-132 effects on Tau acetylation (see Fig. 3B, C) are directly mediated by EP300.

Rbfox1, also known as Ataxin-2-binding protein 1 (A2BP1) or FOX1, is an RNA binding protein (RBP) highly expressed in neuronal tissues^53^ that plays a pivotal role in alternative splicing, mRNA stability and translation in the brain^53,54,55^. Rbfox1 was predicted as another highly scored miR-132 target. We observed that miR-132 mimic reduced Rbfox1 at mRNA and protein levels (Fig. 4A, B). Furthermore, the direct binding and reciprocal regulation of Rbfox1 by miR-132 mimic and inhibitor was validated using the luciferase reporters bearing the WT and mutant Rbfox1 3’UTR, as described above for the GSK3β (Fig. 4C, D). We hypothesized that Rbfox1 may regulate Tau mRNA splicing and/or stability. Indeed, silencing of Rbfox1 by RNAi reduced total mRNA and protein levels of Tau in primary neurons (Fig. 4E, F). Using a Cross-linking and ImmunoPrecipitation (iCLIP) approach, we determined that Rbfox1 directly binds to Tau mRNA (Fig. 4G, H), preferentially via the GCAUG motif site found in its coding region (Figure S3). Therefore, while the exact molecular mechanism remains to be established, Rbfox1 appears as a novel RBP that promotes Tau expression. All together, these data strongly suggest that miR-132 directly regulates Rbfox1 and thereby reduces Tau mRNA stability and/or translation.

Finally, we observed that several proteases implicated in Tau cleavage, including Caspases 3 and 7, and Calpain 2, are regulated by miR-132 (Fig. 4I). One of them, Calpain 2, has been predicted as a conserved direct target of miR-132. In addition to containing a strong putative binding site (Fig. 4C), the direct relationship between miR-132 and Calpain 2 mRNA was supported by their inversely correlated expression observed in the dorsolateral prefrontal cortex of a large cohort of MCI and AD patients (n = 538 subjects) [(Fig. 4J shows the re-analyzed mRNA/miRNA dataset described in^22^. qRT-PCR analysis and luciferase reporter assays confirmed, respectively, that transfections of miR-132 mimic reduced Calpain 2 mRNA expression (Fig. 4A), and this effect was mediated by the direct miR-132 binding to the Calpain 2 3’UTR (Fig. 4D). Therefore, miR-132 reduces Calpain 2 mRNA and protein expression and, thereby, may regulate Tau cleavage. Overall, these data indicate that several Tau modifiers, including GSK3β, EP300, Rbfox1, and Calpain 2 are directly regulated by miR-132 and, thus, collectively contribute to Tau homeostasis in neurons. In addition, regulation of Caspases 3/7 cleavage of tau may also be regulated by miR-132 via PTEN/AKT/FOXO3A signaling^18^.

To further investigate whether any of the direct miR-132 targets involved in Tau metabolism play a dominant role in neuroprotection against Aβ, we compared the effects of miR-132 mimic to that of individual siRNAs cognate to their direct targets (GSK3β, EP300, Calpain 2, Rbfox1, and previously validated Foxo3a). To mimic target repression provided by miR-132, siRNA concentrations and transfection conditions were optimized to ensure 100% transfection efficiency in P301S neurons with 35-45% reduction of target mRNA expression (Fig. 5A). Down-regulation of EP300 and GSK3β resulted in a slightly but significantly improved viability of neurons treated with ½t_max_ Aβ, while the individual downregulation of Calpain 2, Rbfox1, and Foxo3a was insufficient to reduce the Aβ toxicity. Transfections of the five siRNAs together enhanced the neuroprotection relative to the effects of individual siRNAs, but still did not reach the level of protection provided by miR-132 (Fig. 5B). These results indicate that the neuroprotective properties of miR-132 are mediated by multiple target genes, rather than a single key target, and perhaps additional genes beyond the 5 studied here.

**Figure 5.**
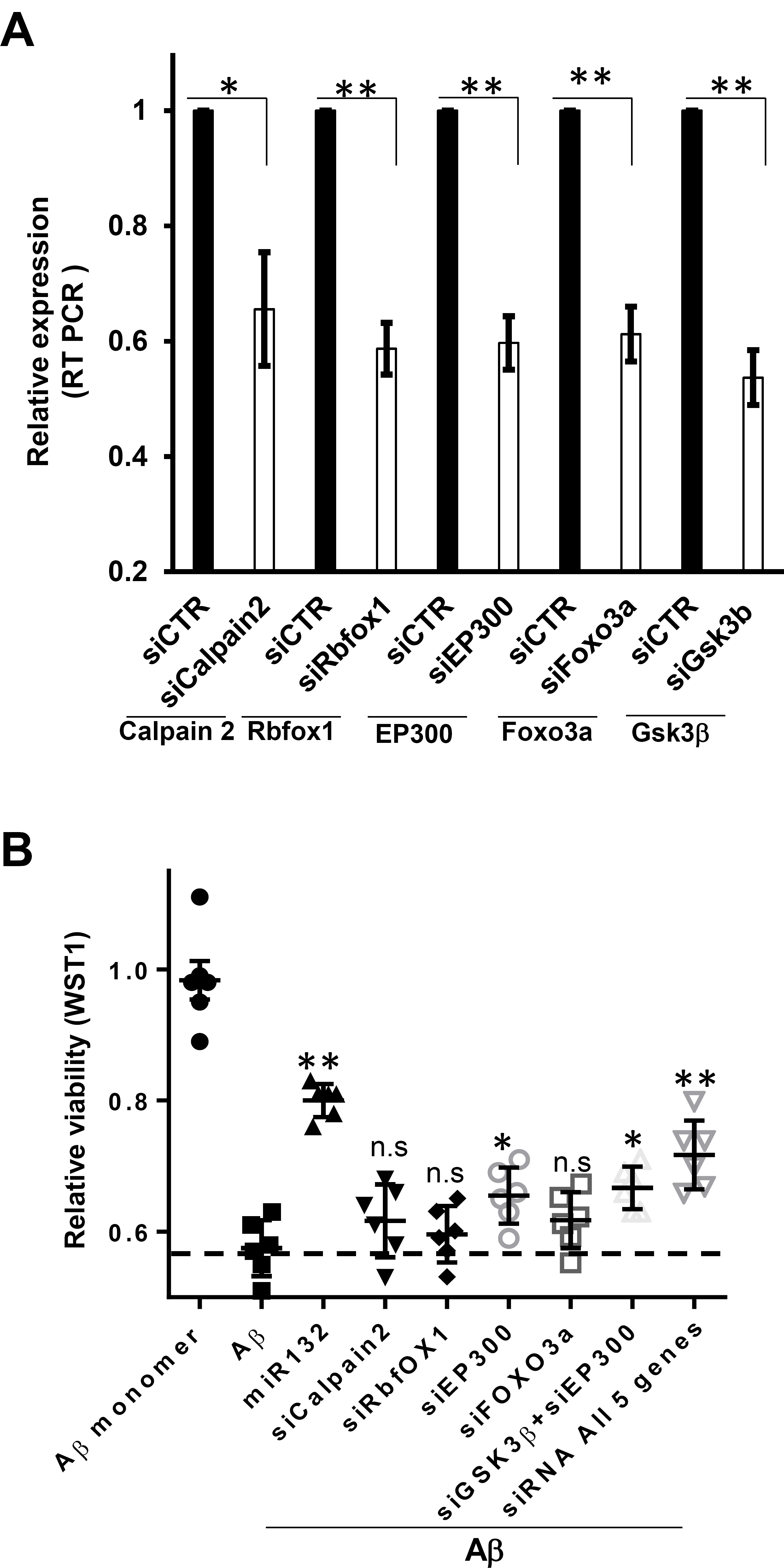
MiR-132 provides stronger neuroprotection than downregulation of its individual targets. **(A)** mRNA levels of Foxo3a, EP300, Calpain2, Rbfox1 and GSK3β mRNAs in primary neurons transfected with cognate siRNAs. (**B**) Relative viability of primary PS19 neurons transfected with miR-132 mimic or individual siRNAs to Foxo3a, EP300, Calpain2, RBfox1, Gsk3β, or siRNAs to all five genes, and exposed to toxic Aβ. *P <0.05, **P <0.005, n = 6. Graphical data are shown as mean +/- SEM.

### Overexpression of miR-132 in PS19 mice reduces caspase-3 activation, Tau hyper-phosphorylation, and neuron loss

Neurons in AD exhibit a ~2-3-fold downregulation of miR-132 levels^18,24^ and similar downregulation was observed in the hippocampus of the 6-months old PS19 mice (Fig. 6A). To investigate the neuroprotective properties of miR-132 *in vivo*, we produced a lentivirus expressing the mature miR-132 from the Synapsin promoter (LV-miR132), as well as a control virus lacking the miR-132 gene (Figure S4). The conditions for durable and spatially defined miR-132 overexpression were optimized for the titrated LV-miR132 injected stereotactically into the hippocampal CA1 area of six-month-old wild type mice (Fig. 6B). The levels of miR-132 overexpression were assessed in the CA1 and the adjacent CA2/CA3 at days 5, 14, and 31 post-injection (Fig. 6C). To reduce non-specific effects and avoid saturation of the system, virus titers of 10^6^ transducing units (TU)/ml provided stable ~2.5-3 fold miR-132 replacement/overexpression in CA1 and ~1.5-2 fold in CA2/CA3 neurons for at least one month were selected for the subsequent experiments (Fig. 6C). This increase in miR-132 expression led to a corresponding downregulation of validated miR-132 targets, including pro-apoptotic FOXO3a, EP300 and downstream effector Bim, and newly established targets GSK3β, Calpain 2 and Rbfox1 (Fig. 6D). These data indicate that this LV system allows sustained functional miR-132 overexpression in mouse hippocampus.

**Figure 6.**
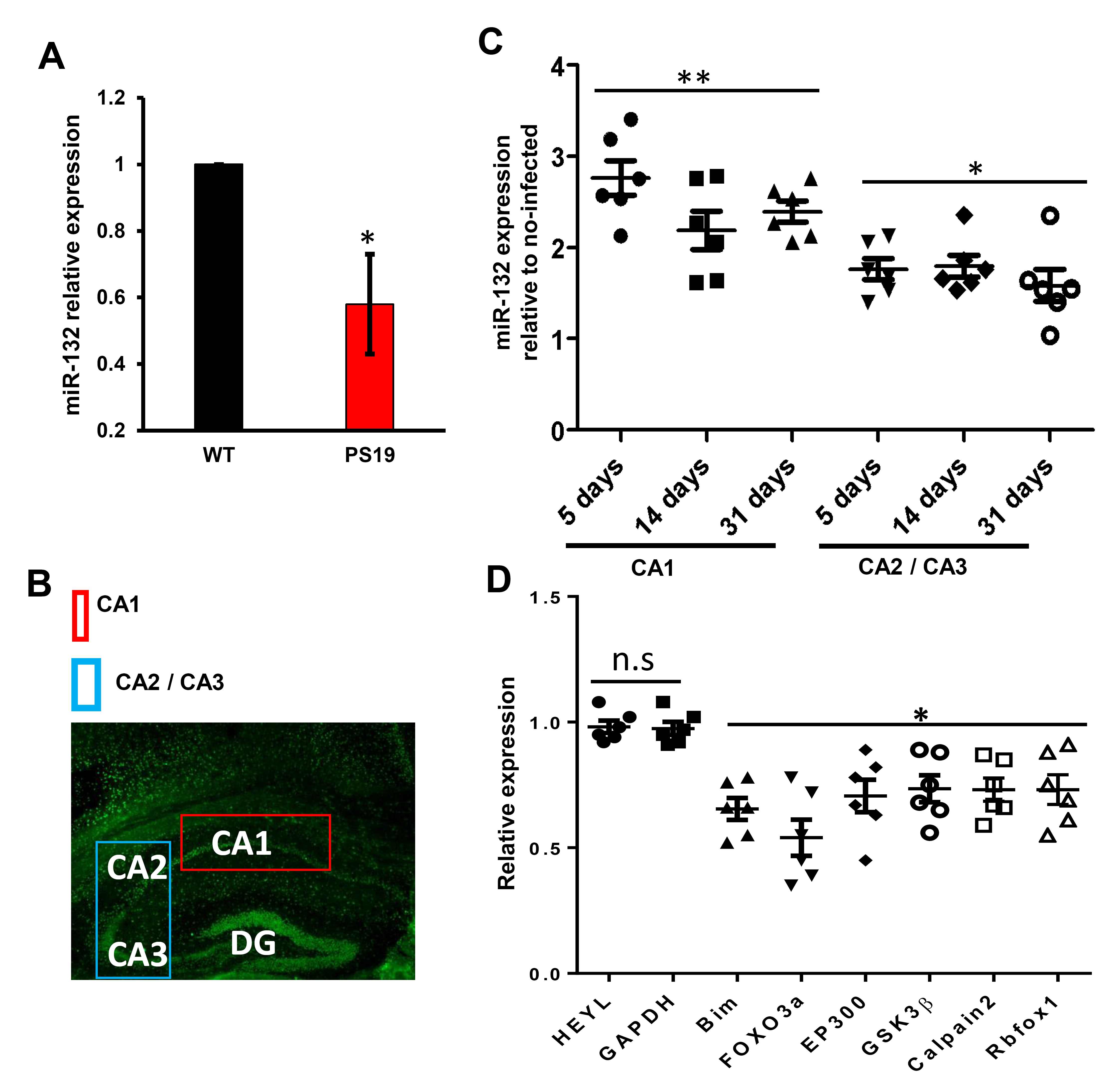
Stable supplementation of miR-132 and downregulation of its targets in the murine hippocampus. **(A)** qRT-PCR analysis demonstrates reduced expression of miR-132 in the PS19 CA1 versus its expression in the CA1 area of 6 months old WT mice. (**B**) NeuN IHC of the mouse hippocampal CA1 area, injected with the miR-132-expressing lentivirus (6.10^6^ TU/ml), and the adjacent CA2/CA3 regions. The boxes depict the areas microdissected for the analysis. (**C**) qRT-PCR analysis demonstrates elevated miR-132 expression in the CA1 and CA2/CA3 regions at days 5, 14, and 31 post-injection of the LV-miR132, relative to the EV (n = 6). (**D**) qRT-PCR analysis shows the reduced mRNA expression of miR-132 targets Foxo3a, EP300, p250GAP, Calpain2, RBfox1, Gsk3β, and downstream Bim, but not of control genes HEYL and GAPDH, in the CA1, at day 31 post-injections. *P <0.01, **P <0.005, n = 6. Graphical data are shown as mean +/- SEM. See also Figure S4.

We then investigated the neuroprotective effects of LV-miR132 in PS19 mice, which develop NFT-like inclusions in the brain and spinal cord starting at around six months of age and evince neuronal loss and brain atrophy by eight months^56^. Experiments were performed to investigate whether ectopic miR-132 expression could prevent or halt neurodengeneration. In the first set of experiments, the viruses were stereotactically injected into the CA1 hippocampal area twice, first at 7.5 months, and then at 9 months of age (Fig.7A). Three experimental groups that included untreated animals, and mice injected to the right hippocampus with either empty LV, or LV-miR132, were analyzed in parallel (15 mice per group). The animals were sacrificed at 10.5 months, and the brains analyzed by immunohistochemistry (IHC) and quantitative image analysis for key markers, including the PHF-Tau, NeuN, activated Caspase-3, and GFAP (Fig. 7B-G). There were ~19% fewer NeuN-positive neurons in the hippocampal CA1 areas and adjacent cortical layers of PS19 mice than in the WT brains (Fig. 7B, D). Cleaved and activated caspase-3 (a marker of apoptosis) was essentially evident in all PS19 brain sections, but absent in WT brains. Similarly, PHF-Tau, a marker of pathology was readily observed in the PS19, but not WT brains (Fig. 7B, F and Figure S5). LV-miR132 significantly reduced the numbers of cells positive for PHF-Tau and cleaved caspase-3 in comparison to the contralateral hemispheres, and to the brains injected with the empty virus, or left untreated (Fig. 7B, E, F). Correspondingly, image quantification indicated that LV-miR132 increased the number of NeuN-positive hippocampal neurons in PS19 mice (Fig. 7D), but GFAP staining was not different between the PS19 groups (Fig. 7G). Consistent with the reduced Tau pathology and neuronal apoptosis, miR-132 increased hippocampal volume relative to the empty LV group (Figure S6). These results demonstrate significant *in vivo* neuroprotection provided by miR-132 in PS19 mice in the progressive stage of neurodegeneration.

**Figure 7.**
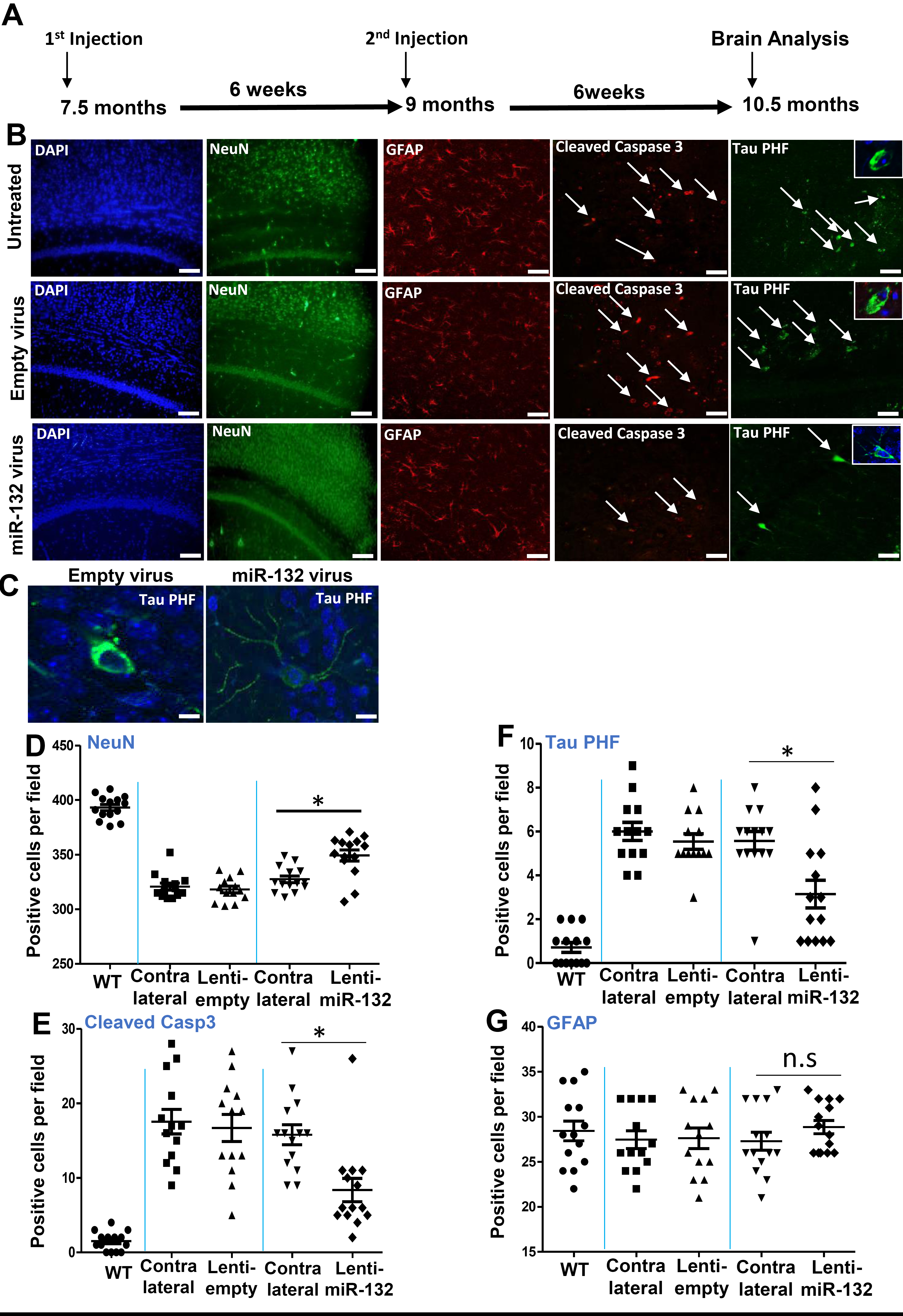
miR-132 supplementation reduces Tau pathology and neuronal loss in PS19 mice. **(A)** Timeline of the LV-miR132 injections and analysis of PS19 brains. (**B**) Representative IHC images of CA1 and adjacent cortical layers stained for NeuN, cleaved Caspase-3, PHF-Tau, glial fibrillary acidic protein (GFAP), and DAPI staining. The cells positive for the activated caspase-3 and PHF-Tau are marked by arrows. Scale bar = 100 μm. (**C**) The tangle-like PHF-positive inclusions (in green) observed in the cell bodies of untreated/EV-treated neurons, and the less intense and more evenly distributed PHF staining in the soma and neurites of apparently intact neurons from miR-132-treated mice. Scale bar = 100μm. (**D** - **G**) Image J quantification of the marker-positive cells per field in the CA1 area for (**D**) NeuN, (**E**) cleaved caspase-3, (**F**) PHF-Tau, and (**G**) GFAP. For all images and quantifications, n = 14 mice per condition, 15 sections per brain. All graphical data are shown as mean +/- SEM, Student’s t-test, *P<0.005. See also Figures S5-S7.

In an additional set of experiments, PS19 mice were stereotactically injected with LV-miR132 a total of three times (i.e. at 3, 4.5, and 6 months, 7 mice per group), before onset of pathology, and analyzed at a time point when pathology is obvious in untreated PS19 mice (i.e. 10 months). In accord with our “treatment” study, supplementation of miR-132 prevented neuronal loss and accumulation of PHF-Tau when administered prior to the emergence of tau pathology (Figure S7). Thus, dependent on the time of administration, overexpression of neuronal miR-132 in the PS19 mice can act both to prevent and ameliorate Tau pathology and neurodegeneration.

### Overexpression of miR-132 enhances hippocampal LTP in WT mice and restores it in PS19 mice

To further investigate functional effects of miR-132 supplementation in the brains of PS19 mice, we measured hippocampal long-term potentiation (LTP) an electrophysiology correlate of learning and memory^57^. Standard high frequency stimulation (HFS) was applied to the CA1 hippocampal region of WT and PS19 mice, and average percent changes in fEPSP slopes relative to baseline stimulation were plotted over time post-HFS (n = 7–15) (Fig. 8A, B). As expected for mice exhibiting neuronal loss^30^ LTP was consistently lower in brain slices from 7.5-month PS19 mice than in age-matched WT mice (136±4%, *n*=15 vs. 165±8%, *n*=12, P<0.001; Fig. 8A]). In WT mice, LTP was enhanced in animals injected with LV-miR132 more than in those injected with empty LV (212±10%, *n*=7 vs. 161±7%, *n*=7, P<0.001; Fig. 8B). In PS19 mice, LV-miR132 treatment dramatically increased LTP to levels greater or comparable to that of untreated WT mice (181±8%, *n*=7 vs. 135±5%, *n*=6, P<0.001; Fig. 8C, D, E). LV-miR132 treatment also potentiated the effects of weak HFS that was insufficient to induce significant LTP in hippocampal slices (115 ± 3% vs. 138 ± 4%, P <0.001; Fig. 8F).

**Figure 8.**
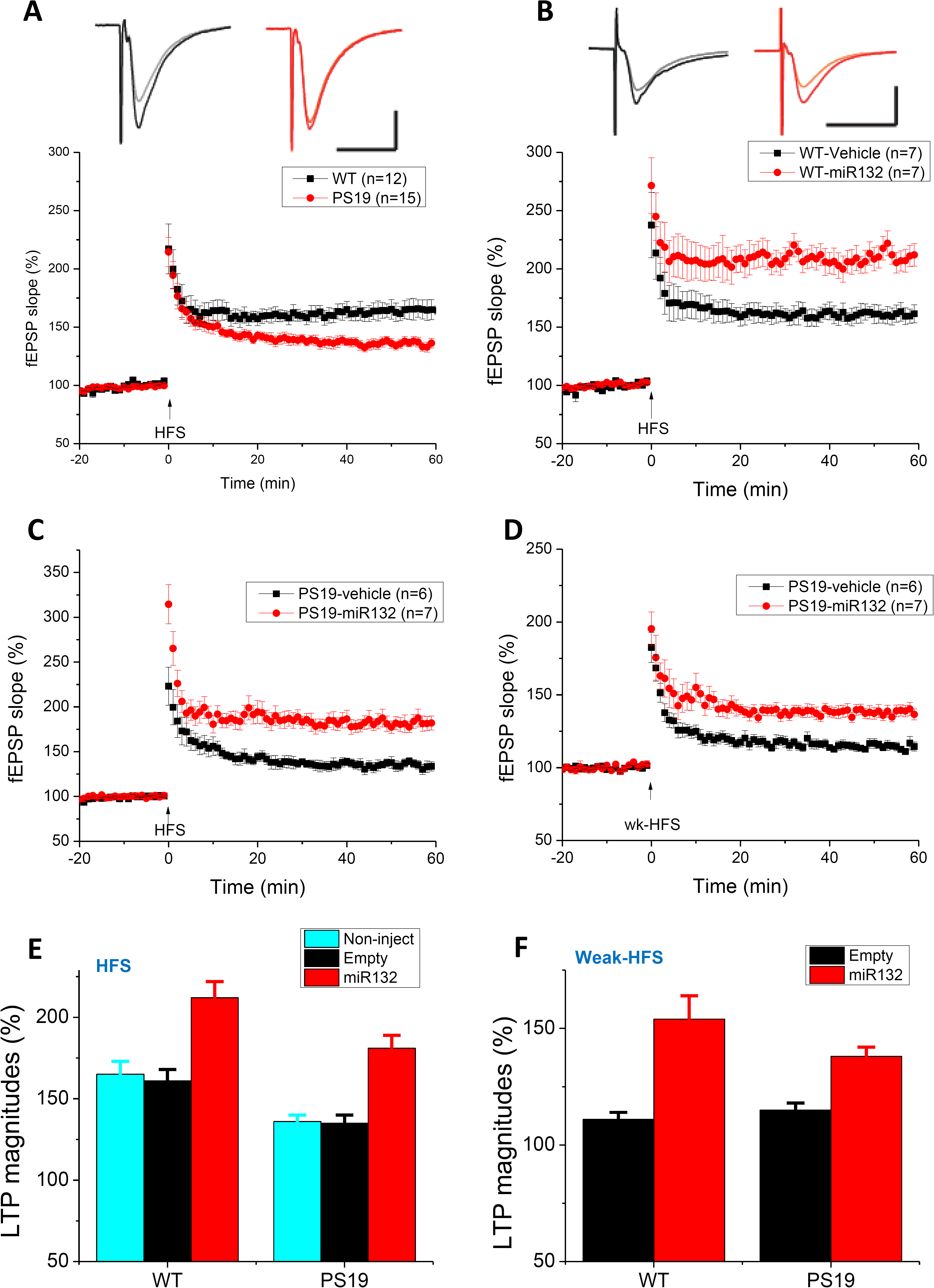
Hippocampal recording demonstrates that miR-132 supplementation rescues LTP impairment in the PS19 mice. (A) The percentage of potentiation of field EPSPs recorded before and after tetanic stimulation of Schaffer’s collaterals in brain slices of PS19 transgenic mice (red) and littermate wild-type mice (black). Each data point shown is the mean ± S.E.M. of results from 8 individual mice of each genotype. (B) Hippocampal miR-132 injection significantly increased LTP induced by high frequency stimulation (HFS, arrow) in the CA1 region of hippocampal slices (red circles, n=7 slices/from 5 mice) vs. those of mice in vehicle (black squares, n=7/4). (C) The reversing effect of miR132 on LTP induced by high frequency stimulation (HFS, 100 Hz last 1 sec, 2 trains separated by 20 sec) in PS 19 mice injected with either empty lentiviruses (black squares, n=6/4) or LV-miR132 (red circles, n=7/4). (D) The miR132 injection enable a weak HFS (100 Hz last 1 sec) induced LTP in PS19 mice injected with LV-miR132 (red circles, n=7/5) not the empty lentiviruses (black squares, n=6/4). Bar diagram summarizing the LTP experiments with WT and PS19 mice induced by standard HFS (E) or weak HFS (F). Inset traces are typical field excitatory postsynaptic potentials (fEPSPs) recorded before (gray or orange) and after (black or red) HFS for each condition. Horizontal calibration bars: 10 ms; vertical bars: 0.5 mV.

## Discussion

Multiple lines of evidence support the significance of reduced miR-132 activity in AD and related neurodegenerative conditions. First, from many independent attempts to define miRNAs linked to AD pathology miR-132 has emerged as the top molecule significantly associated with both plaques and tangles in a variety of disease affected brain areas^18,19,20,21,22,36,58^. miR-132 is downregulated starting at Braak III stage, before neuron loss, and miR-132 reduction is evident in phospho-tau positive neurons^19,24^. Furthermore, miR-132 downregulation has been described in other neurodegenerative disorders linked to aggregation and accumulation of misfolded protein Tau, including frontotemporal lobar degeneration and progressive supranuclear palsy^58,59,60^. Although miR-132 downregulation in the latter classes of tauopathies still requires validation in larger brain cohorts, the data suggest a possible common mechanism underlying miR-132 dysregulation in both AD and primary tauopathies. Second, miR-132 knock-out impairs memory formation and retention in adult mice^21^, induces Tau aggregation, and aggravates both tau and amyloid pathologies in transgenic mouse models^19,20,21^. Third, our high-content screen for miRNA modulators of neuroprotection against Aβ and glutamate excitotoxicity performed in this study, identified miR-132 as the top hit (Fig. 1). Finally, miR-132 has important regulatory functions in neuron development, synaptic plasticity, and survival^61^. Of note, additional neuroprotective (e.g., miR-29 and miR-129) and “neurotoxic” (e.g., miR-26b and miR-34a) miRNAs identified in our screen have been previously implicated in the regulation of critical genes and pathways in AD^22,62,63^; and it will be important to investigate these hits in future studies.

Several targets and signaling pathways may underlie miR-132 neuroprotective functions. Some of them, such as p250GAP, RASA1, and MeCP2, mediate the role of miR-132 in neurite extension, arborization and synaptogenesis^64^. Other targets, such as PTEN, p300, FOXO3a counteract AKT pro-survival signaling; their derepression observed in AD neurons and probably caused by miR-132 downregulation may induce expression of the key apoptotic effectors Bim and Puma, leading to activation of caspases and apoptotic signaling^18^, and also promoting Tau cleavage. Furthermore, recent reports suggest certain direct miR-132 targets implicated in Aβ and Tau metabolism^61^, including the Tau mRNA itself^20^. However, miR-132 does not appear to regulate Tau in human neurons directly, and the miR-132 binding site is not conserved in primate Tau mRNA ^36^. Tau homeostasis is tightly controlled at multiple levels, and we report here that miR-132 regulates several key factors affecting tau production, posttranslational modifications, and proteolysis (Fig. 3). Specifically, miR-132 regulates tau phosphorylation (via direct targeting of GSK3β), acetylation (via a EP300), and cleavage (through calpain 2 and caspases-3/7), and it also reduces Tau mRNA via the direct targeting of the RNA-binding protein, RBFOX1. Tau hyperphosphorylation at Ser-396/404 (PHF-1 epitope), largely mediated by GSK3β, affects microtubule dynamics and NFT accumulation, which is considered a hallmark cytopathology in AD and other tauopathies^65^. Since we validated both major tau kinase GSK3β and acetylase EP300 as the direct miR-132 targets (Fig. 4 and^18^, and additional tau kinase CDK5 is also indirectly repressed by miR-132 via NOS1 signaling^36^, thus miR-132 emerges as the major regulator of the posttranslational modifications of Tau.

Our work also demonstrates that miR-132 regulates Tau cleavage and implicates a newly validated target calpain 2, as well as caspases 3 and 7, in this event. Tau is cleaved by multiple proteolytic enzymes which facilitate its degradation and clearance. However, if allowed to accumulate, some of these fragments become aggregated and/or hyperphosphorylated, and neurotoxic^34,35,36^. For instance, Tau cleavage by calpain 2 produces a 17 kDa neurotoxic fragment, and significant amounts of this fragment are found in the brains of patients with tauopathy^39,40^. Mutations in calpain in transgenic flies were shown to prevent Tau toxicity^40^. In addition to the proteolysis of Tau, Calpain also cleaves p35, the principal activator of Cdk5, into p25, which results in the hyperactivation of both Cdk5 and GSK3β, and thereby induces tau hyperphosphorylation^66,67^. Here we demonstrate that miR-132 not only regulates Calpain 2 directly, but its levels also inversely correlate with the levels of Calpain 2 mRNA in hundreds of AD brains, suggesting that miR-132 is a primary regulator of Calpain 2 expression in the brain, responsible for its upregulation in AD. In addition, caspase 3/7 activity, modulated by miR-132 indirectly (Fig. 4I), likely through PTEN/FOXO3/Bim signaling^18^, may also contribute to Tau cleavage and the observed release of tau fragments from neurons (Fig. 3D, E).

We also validated Rbfox1, an RNA-binding protein highly expressed in neuronal tissues^53^, as another direct miR-132 target (Fig. 4B-D). By binding to the GCAUG element, Rbfox1 plays a pivotal role in alternative splicing^53^, mRNA stability^54,55^ and translation^54^. The Rbfox1 knockout mice have a significant increase in neuronal excitability in the dentate gyrus^68^, and a recent study identified a link between Rbfox1 protein and AD^55^. We demonstrate that Rbfox1 binds to and stabilizes neuronal Tau mRNA (Fig. 4E-H). Altogether, considering the additional miR-132 target PTBP2 previously implicated in Tau mRNA splicing^20^, these data position miR-132 as the principal regulator of various aspects of Tau homeostasis and suggest a mechanistic link between the miR-132 downregulation and Tau pathology observed in disease.

Overall, our work supports miR-132 as the master regulator of neuronal health. In addition to its distinct functions in synaptogenesis, neuronal activity, plasticity, memory, and neuronal viability^64,69^, miR-132 appears to regulate Tau metabolism, and its downregulation in AD and other neurodegenerative diseases likely promotes pathogenesis by perturbing multiple signaling pathways. Interestingly, an initial increase in miR-132 levels during early AD Braak stages I–II in the human prefrontal cortex has been described, which contrasts with the decrease seen at more advanced stages of the disease^24^. A similar bi-phasic miR-132 expression pattern has been reported in prion disease^70^, suggesting that miR-132 is part of an initial neuroprotective response. Subsequent downregulation, however, can aggravate the effects of Aβ and Tau toxicities^21^. Of note, miR-132 is regulated by the activity-dependent cAMP-response element binding (CREB) transcription factor, and its expression pattern in the AD brain mimics that of the brain-derived neurotrophic factor (BDNF)^14^. Also, alterations in DNA methylation that affect gene expression and perhaps the onset of AD^71^, may play a role in miR-132 downregulation in neurons, as demonstrated for some cancer cells ^72,73^.

Collectively, these data suggest that miR-132 replacement or normalization in tauopathies, such as AD, may provide a much-desired neuroprotective effect. Lowering Tau alone with antisense oligonucleotides (ASO) has recently been shown as therapeutically beneficial for tauopathies ^74^. Here we provide a proof-of-principle for miR-132 replacement as a novel neuroprotective strategy to reduce Tau pathology and simultaneously promote nerve growth and regeneration, enhance neuronal survival, and consequently improve cognition. miR-132 supplementation protects against strong toxic stimuli even in highly vulnerable and damaged Tau-mutant neurons, *in vitro* and *in vivo*. It had preventive effects in young presymptomatic PS19 mice, and reduced neuronal loss and Tau pathology even when pathology was already established. The PS19 model exhibits broad brain and spinal cord pathology resulting in severe cognitive, motor, and visual impairments^30,75^. Since our proof-of-concept study relies on the lentivirus-mediated local unilateral miR-132 supplementation to the CA1 region, it did not allow examination of the effects on global readouts such as neurologic and behavioral phenotypes. To overcome this limitation, broader distribution of miR-132 mimics in the CNS will be required.

Notably, small molecules and other types of inhibitors of major miR-132 targets have entered clinical trials. These include inhibitors of GSK3β kinase, EP300 acetylase, calpains, as well as Tau-targeting antibodies and ASO drugs^76,77,78^, the latter was presented at the 142nd Annual Meeting of the American Neurological Association, October 2017]. Remarkably, miR-132 is the natural inhibitor of each of these factors. Therefore, its replacement can provide a multi-hit approach and ensure the benefits of combination therapies. MiR-132 replacement strategies for tauopathies will largely rely on the development of miRNA-mimicking oligonucleotides and technologies for their delivery to the brain, and leverage recent advances in the field of oligotherapeutics. Notably, the first “breakthrough” oligonucleotide-based drug for a neurologic disease has recently gained fast FDA approval^79^, and many more are at different stages of clinical development for a wide spectrum of disorders (including AD, amyotrophic lateral sclerosis, Huntington’s disease, and FTD). AD and other tauopathies have so far proven refractory to small molecules and biological drugs, and miRNA mimics emerge as a new and promising class of therapeutics. Our work validates miR-132 as a first-line candidate for development of such neurotherapies.

## Acknowledgment

We thank Dr. Li Gan for providing antibodies for acetylated Tau, Drs. Andy Billinton and Mike Perkington (MedImmune) for the gift of TauAB antibody, and Jie Shen for valuable discussions and equipment access. We thank members and advisers of the Tau Consortium for valuable discussions. This work was supported by grants from Alzheimer’s Association (NIRG-09-132844), and Tau Consortium/ Rainwater foundation.

## Conflict of interest

None.

## Author Contributions

R.E.F and A.M.K conceived the project and analyzed the data, R.E.F performed most experiments, S.L., Z.C., T.M., S.G., Z.W., D.T.B., R.R., A.C., and A.E. assisted with experiments, D.J.S., K.C.S., and D.M.W. contributed to data analysis, and R.E.F and A.M.K. wrote the manuscript. All authors critically reviewed the manuscript.

## Methods

### Primary Neuronal Cultures and their analysis

Primary cortical and hippocampal neuron cultures were prepared from WT E18 and PS19 mice (RRID:IMSR_JAX:008169) and human fetal cortical specimens (provided by Advanced Bioscience Resources). All studies have been approved and performed in accordance with Harvard Medical Area and BWH Standing Committee (IACUC) guidelines. Brain tissues were dissected, dissociated enzymatically by papain, and mechanically by trituration through Pasteur pipette, plated and cultured as previously described^36^. Imaging of the cultures was performed using the IncuCyte^TM^ Live-Cell Imaging System (Essen BioScience). Cell confluency, cell body number, neurite length and branching points were monitored and quantified using IncuCyte^TM^ software. Neuron viability was measured using the WST1 assay, following the manufacturer’s instructions (Roche).

### Transfections of primary neurons

Transfections of miRNA mimics (Miridian oligos at 20nM final concentration, Dharmacon), inhibitors (LNA-containing at 50nM, Exiqon), siRNAs (at 25nM), and the corresponding control oligonucleotides of the same chemistries, to primary mouse and human neurons were carried out using the NeuroMag technology (OZ Biosciences). The oligonucleotides were mixed with the nanoparticles in Neurobasal medium, at room temperature for 15 min. The transfection complexes were then added to the cultured neurons and placed on a magnetic plate at room temperature for additional 15 min. The cultures were further incubated with the transfection mixture in a standard culture incubator overnight. Half of the media was replaced next morning, and the remaining media was replaced at later time points.

### SEC-isolation of Aβ monomer and preparation of ½-*t*_max_ Aβ_*(1–42)*_

Based on the general consensus that aggregation of Aβ is required for toxicity, we employed a partially aggregated preparation of Aβ(1–42) that contained both amyloid fibrils and Aβ monomer^18,26,27^. This preparation is referred to as 1/2*t*-max because it is produced by incubating Aβ monomer for a period that yields half the maximal level of thioflavin T. When used at concentration ≥ 10μM 1/2*t*max can cause the compromise and death of cultured rodent and human neurons within a period of a few days ^18,26,27^. Synthetic monomer Aβ_(1–42)_ (human sequence) was obtained from rPeptide (A-1165-2). Briefly, Aβ(1–42) was dissolved at 1 mg/ml in 50 mM Tris–HCl, pH 8.5, containing 7M guanidinium HCl and 5 mM ethylenediamine tetraacetic acid, and incubated at room temperature overnight. The sample was then centrifuged at 16,000 × *g* for 30 minutes and the upper 90% of supernatant applied to a Superdex 75 10/300 size exclusion column (GE Healthcare Biosciences), eluted at 0.5 ml/minute with 50 mM ammonium bicarbonate, pH 8.5. Absorbance was monitored at 280 nm. Fractions of 0.5 ml were collected. Peak fractions were pooled and the concentration of Aβ determined using ε_275_ = 1,361/M/cm^80^. Thereafter ½-*t*_max_ Aβ_(1–42)_ was prepared as described previously^81^. Aβ _(1–42)_ synthetic monomer was resuspended in 10.9 mM HEPES, pH 7.8, and filtered through a 0.22 μm filter (Millipore) in a sterile hood. An aliquot of the filtered sample was used for concentration determination by measuring the absorbance at 275 nm, using the extinction coefficient 1361 M^−1^ cm^−1^ in a spectrophotometer with a 1 cm path length microcuvette. The remainder of the sample was diluted with sterile buffer to 80 μM and all further manipulations were performed on ice. Thioflavin T (ThT) was added to a portion of the stock Aβ solution and was incubated with shaking (700 rpm) at room temperature and fluorescence was monitored at 20 min intervals to determine the interval required to reach ½-*t*_max_. The remainder of the Aβ stock solution that did not contain ThT was incubated in the same conditions and the reaction was stopped at 180 min, according to the previously established curve, with additional replicates containing ThT to confirm reproducibility of the aggregation kinetics. Aβ ½*t*_max_ samples were aliquoted on ice, flash frozen on dry ice, and stored at −80°C.

### Real-time quantitative RT-PCR

Total RNA from primary cultures and mouse brains was extracted with Exiqon RNA isolation kit, according to the manufacturer’s instructions. For miRNA quantifications, TaqMan^®^ miRNA assays (Life Technologies) were used, following the manufacturer’s protocol. Relative mRNA levels were monitored by qRT-PCR reactions with specific primers (listed in the supplemental table S1), on the ViiA-7 System (Thermo Fisher Scientific). Threshold cycles (Cts) were generated automatically, and the relative expressions were shown as 2^−∆Ct^. Relative mRNA levels of miR-132 target genes were normalized to the geometrical mean of 18 rRNA, ACTB and PABP2 mRNAs. Relative miRNA levels were normalized to the geometrical mean of the uniformly expressed miR-99a, miR-181a, and U6 snRNA.

### Western blotting analysis

Proteins have been extracted and the concentrations determined by Pierce^TM^ BCA Protein Assay Kit. For Western blot analysis, the proteins have been resolved on the SDS-PAGE, transferred to 0.45 μm nitrocellulose membranes (BioRad), blocked with 5% non-fat dry milk in PBS with 0.1% Tween 20, and processed for immuno-detection. The following primary antibodies were used following manufacturer’s instructions: Tau 5, Tau 46, Tau PHF, for Rbfox1, Calpain 2, cleaved Caspase-3, cleaved Caspase-7, GSK3β, EP300, and β-Actin (all from Cell Signaling). Anti-acetyl-Tau AC312 (rabbit anti-ac-K174 Tau) and MAB359 (rabbit anti-ac-K274 Tau) kindly provided by Li Gan’s laboratory were used at 1/5,000 dilution. Antibody detection was performed with the HRP-coupled goat secondary anti-mouse or anti-rabbit antibodies (Immunoresearch), followed by the ECL reaction (Perkin Elmer) and exposure to Fuji X-ray films. The films were scanned and signals quantified and analyzed using the ImageJ software.

### Detection of intracellular and extracellular Tau using ELISAs

Two ELISAs were used in this study. One which is highly similar to clinically approved assays which employ mid-region directed mAbs and are often erroneously refered to as total tau assays, and the other a novel C-terminal ELISA that uses mAbs specific for the C-terminus and MTBR domains of Tau. The mid-region tau ELISA was performed essentially as described previously^44^. Briefly, black half-area high binding 96-well plates (Greiner Bio-One, Frickenhausen, Germany) were coated with the anti-tau antibody, BT2, at 2.5 μg/ml in TBS, pH 7.4, with 25 μl added to each well and incubated at 37°C for 1 h. Wells were washed 3 times with TBS containing 0.05% Tween 20 (TBS-T) and then blocked with 3% (w/v) BSA in TBS for 2 h with shaking on an orbital shaker (Woodbridge, NJ, USA) at 300 rpm. Thereafter, wells were washed 3 times with TBS-T. Samples were analyzed in duplicate (25 μl/well), whereas blanks and Tau441 standards (7.8–8000 pg/ml) were analyzed in triplicate. Standards and samples were diluted with assay buffer and incubated overnight at 4°C. The next day, plates were shaken on an orbital shaker for 1 h to allow the assay to return to room temperature. Subsequently, 25 μl of Tau5-alkaline phosphatase in TBS-T containing 1% (w/v) BSA was added to each well and incubated for 1 h with shaking on an oribtal shaker. Tau5 was conjugated to alkaline phosphatase using a Lightning-Link alkaline phosphatase conjugation kit (Innova Biosciences, Babraham, Cambridge, England). Wells were washed five times and 50 μl of chemiluminescent substrate (Tropix CDP-Star Sapphire II; Applied Biosystems, Foster city, CA) was added and incubated with shaking on an orbital shaker for 30 min. Chemiluminescence was measured using a Synery H1 plate reader (Biotek, Winooski, VT). Standard curves were fitted to a five-parameter logistic function with 1/Y2 weighting, using MasterPlex ReaderFit (MiraiBio, San Bruno, CA). The lower limit of quantification (LLOQ) is defined as the lowest standard point with a signal higher than the average signal for the blank plus 9 SDs and a percent recovery ≥100±20%.

The LLOQ was calculated for each plate and for the results shown the LLoQ was 31 pg/ml. The C-terminal ELISA was performed exactly as for the mid-region assays except the polyclonal antibody K9JA (243-441aa) was used for capture and the mAb TauAB (425-441aa) was used for detection. For the results shown the LLoQ of the assay was 7.8pg/ml.

### iCLIP

iCLIP was performed according to the published protocol^82^, with minor modifications. Briefly, mouse neurons were irradiated with UV-C light to covalently cross-link proteins to nucleic acids (400 J/m^2^). Upon cell lysis, samples were subjected to DNase treatment and RNA was partially fragmented using low concentrations of the RNase I (0.002 U/ml, 5 min), following by the treatment with the RNase inhibitor (RNAsin Plus at 0.5 U per μl) to quench RNase activity. The Rbfox1–RNA complexes were immunopurified using the anti-Rbfox1 antibody (Cell Signaling) immobilized on immunoglobulin G–coated magnetic beads. RNA was isolated and precipitated, and the RT-PCR reactions performed with the Tau-specific primers to amplify different segments of the mRNA.

### Validation of miR-132 targets by Luciferase Reporter Assay

Full-length 3′ UTR sequences of Gsk3β, Calpain2 and Rbfox1 were cloned into psiCHECK2 plasmid (Promega, C8021), downstream of *renilla* luciferase, using *XhoI* and *NotI*. Mutations in the miR-132 binding sites were introduced to these constructs using the QuikChange Multi Site-Directed Mutagenesis Kit (Stratagene). Primers used for cloning and mutagenesis are indicated in table S1. Four hundred nanograms of the psiCHECK-based constructs were co-transfected with either miridian miRNA mimics (at 25nM final concentration) or LNA inhibitors (50nM), in Lipofectamine 2000, to the SH-SY5Y cells grown in 96-well plates. Alternatively, to validate the targets in primary neurons, 1μg of psiCHECK2 constructs per well was used in 24-well plates. Two days after transfections, luciferase luminescence was measured using the Dual-Glo Luciferase Assay System (Promega, E2920) and detected with Infinite F200 plate reader (TECAN). Renilla luminescence was normalized with that of firefly and the signals were presented as renilla/firefly relative luminescence.

### Lentivirus production and stereotaxic brain injections

For lentivirus production, the miR-132-expressing PL13-pSyn-mmu-miR-132-IRES2-EGFP, or control PL13-pSyn-IRES2-EGFP plasmid were co-transfected with packaging psPAX2 plasmids and VSV-G envelope-expressing plasmid (Addgene plasmids #12259 and #12260), and the viruses concentrated by additional ultracentrifugation at 25,000 rpm. Lentivirus titers were determined by PCR and functional titer was further determined by serial dilutions in 293T cells, using GFP fluorescence. Positive cells were counted and the titer was estimated using the following formula: titer (TU/mL) = number of transduced cells in day 1 x percentage of fluorescent positive cells x 1,000/volume of lentivirus used (mL). The lentivirus expressing miR-132 (LV-miR132) or empty vector (EV) (2µl) were stereotactically injected at 6 x 10^6^ TU/ml to the CA1 region of the right hippocampus (Bregma coordinates: 2.5 mm posterior, 1.7 mm lateral, and 1.8 mm ventral) P–A, 0.5 mm; C–L, 1.7 mm; and D–V, 2.3 mm) of C57BL/6J and PS19 mice. The animals were randomized to the treatment and control groups. All animal studies have been approved and performed in accordance with Harvard Medical Area and BWH Standing Committee for Animal Care (IACUC) guidelines.

### Immunohistochemistry

Mice were sacrificed by CO2 exposure following cervical dislocation, and the brains fixed in 4% paraformaldehyde, embedded, and cryo-sectioned. The 16-μm-thick sections were immunostained for NeuN, GFAP, Cleaved Caspase-3 and Tau-PFH (with antibodies from Cell Signaling). The sections were first incubated in the blocking solution (7.5% NGS; 0.4% Triton; 1% BSA; PBS) for 2 hr, followed by the overnight incubation in antibody-containing solution (5% NGS; 0.2% Triton; 0.5% BSA; PBS), and 2hr 30min-incubation with a secondary antibody (either AlexaFluor 568 or AlexaFluor 488; Invitrogen). IHC was observed using Zeiss confocal microscope at 20x magnification, and the images were processed with a computerized image analysis system (ZEN 2012 SP2 Software, Zeiss).

### Electrophysiology

Mice were sacrificed by isoflurane anesthesia and brains were quickly removed and submerged in ice-cold oxygenated sucrose-replaced artificial cerebrospinal fluid (ACSF) cutting solution containing 2 KCl, 2 MgSO_4_, 1.25 NaH_2_PO_4_, 1 CaCl_2_, 1 MgCl_2_, 26 NaHCO_3_, 10 D-glucose, pH 7.4, 315 mOsm). Transverse slices (350 μm thick) were cut with a vibroslicer from the middle portion of each hippocampus. After dissection, slices were incubated in ACSF containing (in mM): 124 NaCl, 2 KCl, 2 MgSO_4_, 1.25 NaH_2_PO_4_, 2.5 CaCl_2_, 26 NaHCO_3_, 10 D-glucose, pH 7.4, 310 mOsm, in which they were allowed to recover for at least 90 min before recording. A single slice was then transferred to the recording chamber and submerged beneath continuously perfused ACSF saturated with 95% O_2_ and 5% CO_2_. The slices were incubated in this chamber for 20 min before stimulation at RT (~24°C). Standard field excitatory postsynaptic potentials (fEPSP) were recorded in the CA1 region of the right hippocampus. A bipolar stimulating electrode (FHC, Inc., Bowdoin, ME) was placed in the Schaffer collaterals to deliver test and conditioning stimuli. A borosilicate glass recording electrode filled with ACSF was positioned in stratum radiatum of CA1, 200~300 μm from the stimulating electrode. fEPSP in CA1 were induced by test stimuli at 0.05 Hz with an intensity that elicited a fEPSP amplitude of 40~50% of maximum. Test responses were recorded for 20-30 min before the experiment. To induce LTP, two consecutive trains (1 s) of stimuli at 100 Hz separated by 20 s, a protocol that induces LTP lasting ~1.5 hr in wild-type mice of this genetic background were applied to the slices. The field potentials were amplified 100x using Axon Instruments 200B amplifier and digitized with Digidata 1322A (Axon Instruments). The data were sampled at 10 kHz and filtered at 2 kHz. Traces were obtained and analyzed using the Clampfit 9.2.

## Legends for Supplemental figures

**Figure S1.**
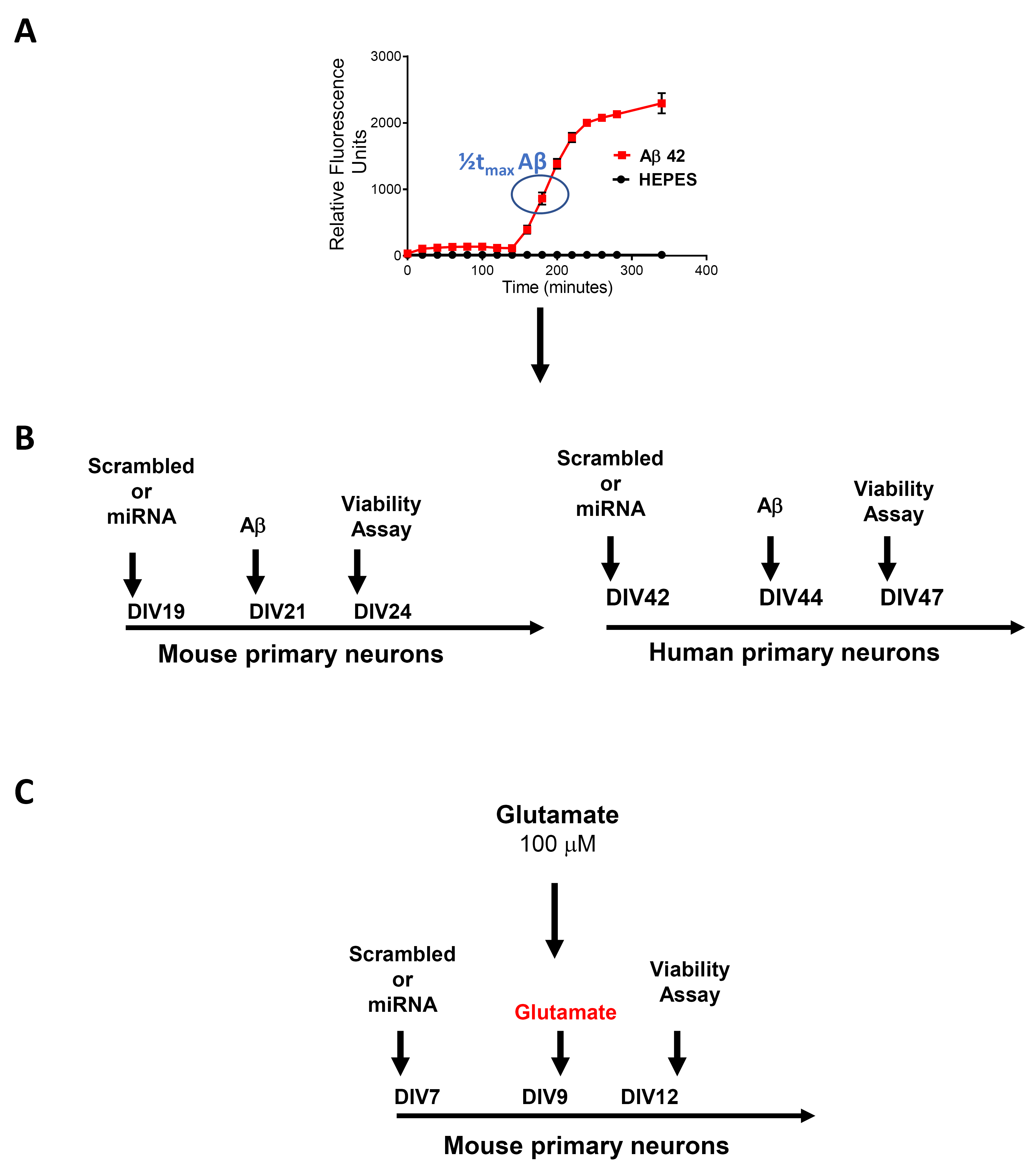
Generation of partly aggregated Aβ and schematics indicating the time course of neuroprotection experiments. **(A)** Aggregation of SEC-isolated Aβ (1–42) was followed using a continuous thioflavin T (ThT)-binding assay and fluorescence values are expressed in relative fluorescence units and plotted versus time. The point at which sample was collected (1/2*t* max) and used for toxicity experiments is indicated with the blue oval. Freshly SEC-isolated, unaggregated monomer (i.e. equivalent to t=0) was used as a control. (**B**) Time course of neuroprotection experiments on mouse and human primary neurons exposed to Aβ or (**C**) excitotoxic glutamate.

**Figure S2.**
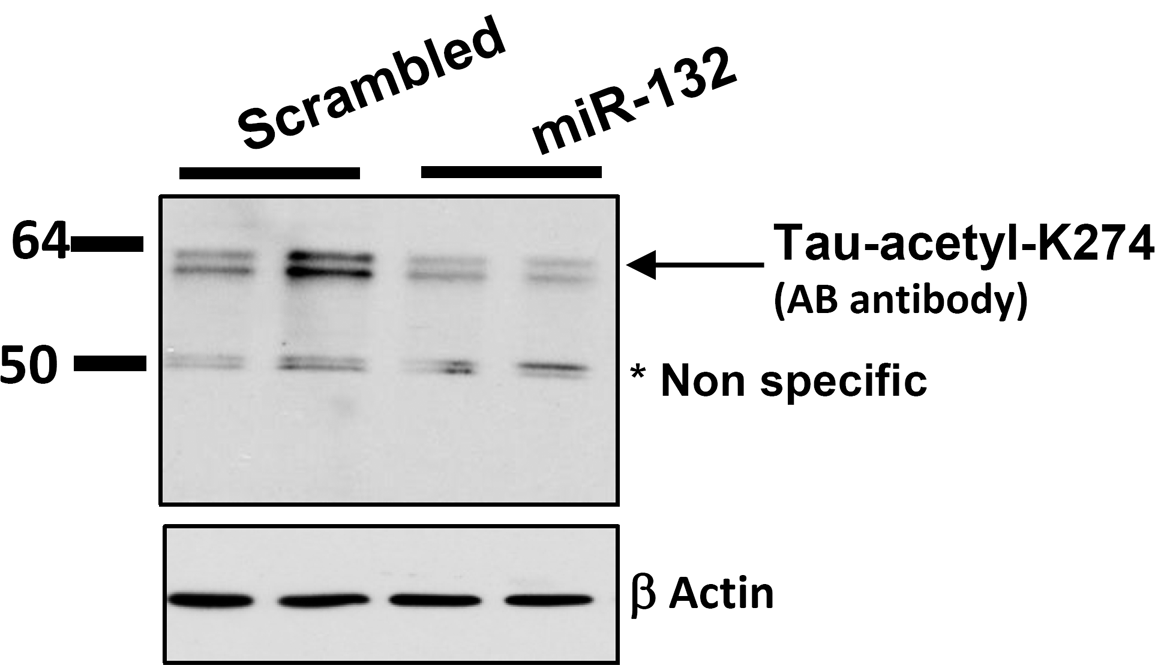
MiR-132 reduces Tau acetylation at K274. Western blot analysis of mouse primary neurons transfected with either miR-132 mimic or scramble oligonucleotide demonstrates that miR-132 reduces Tau acetylation at the position K274.

**Figure S3.**
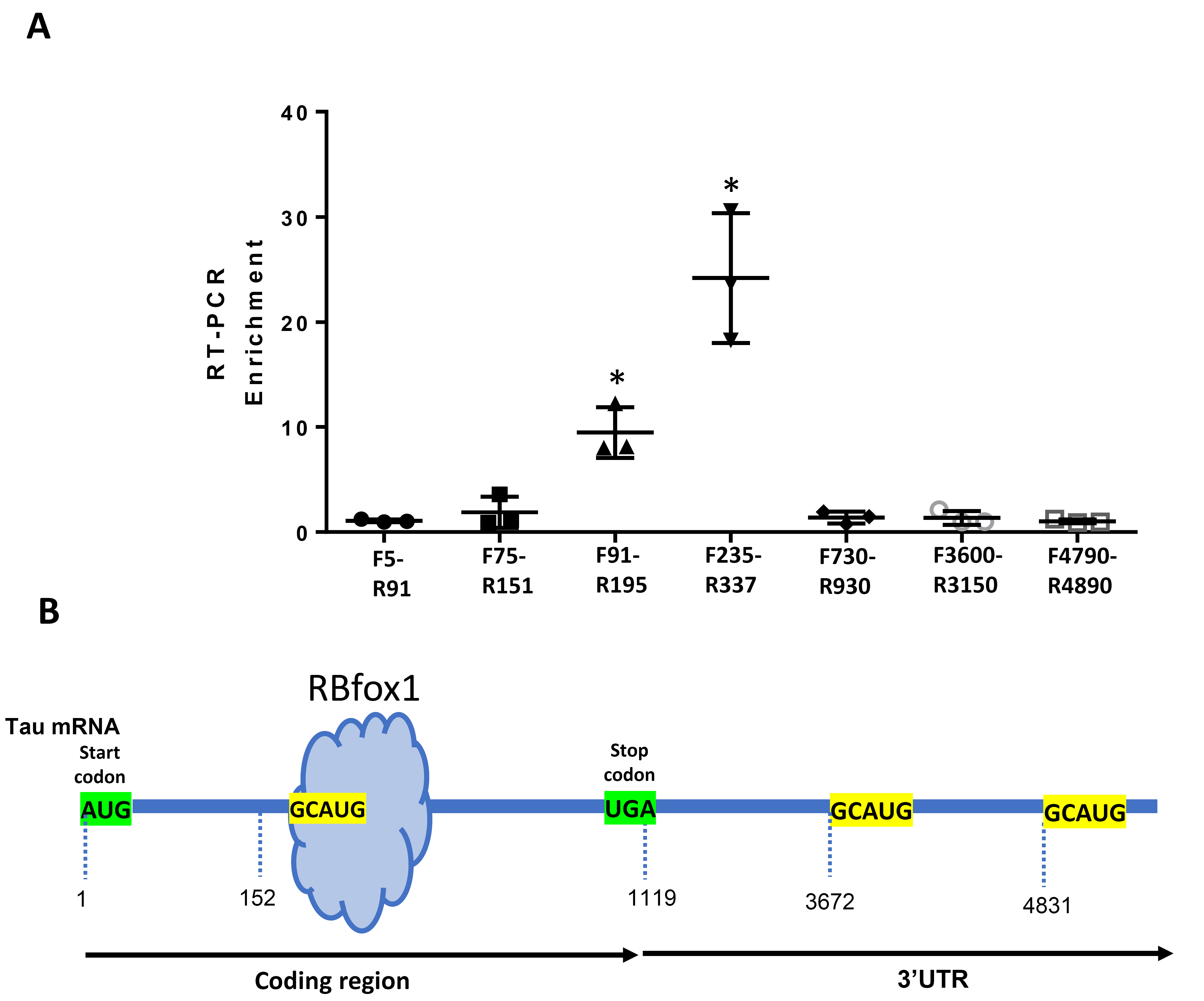
Mapping of the Rbfox1-Tau mRNA interaction in mouse primary neurons. **(A)** RNA immunoprecipitated with the Rbfox1-specific antibody was analyzed by qRT-PCR with primers amplifying various regions of Tau mRNA, including the coding region and 3’UTR. The data are presented as fold enrichment relative to IgG control; mean +/- SEM, n=3, *P<0.005. (**B**) Schematic view of the putative Rbfox1-binding GCAUG motifs in Tau mRNA. In mouse neurons, Rbfox1 binds to the Tau mRNA preferentially via the GCAUG site found in the coding region.

**Figure S4.**
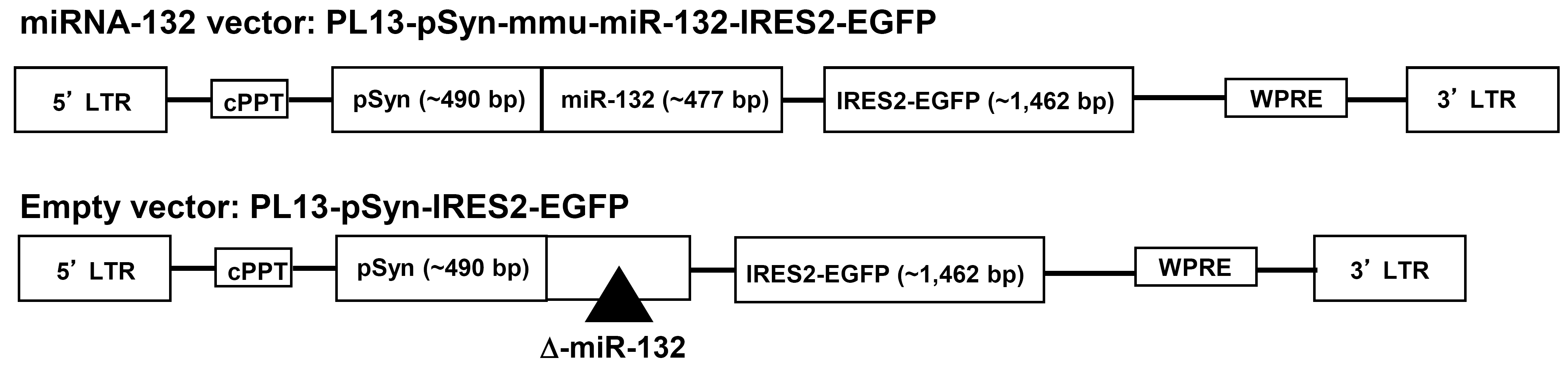
Schematic diagram of constructs used for the lentivirus production of LV-miR132 and the corresponding negative control (Empty vector).

**Figure S5.**
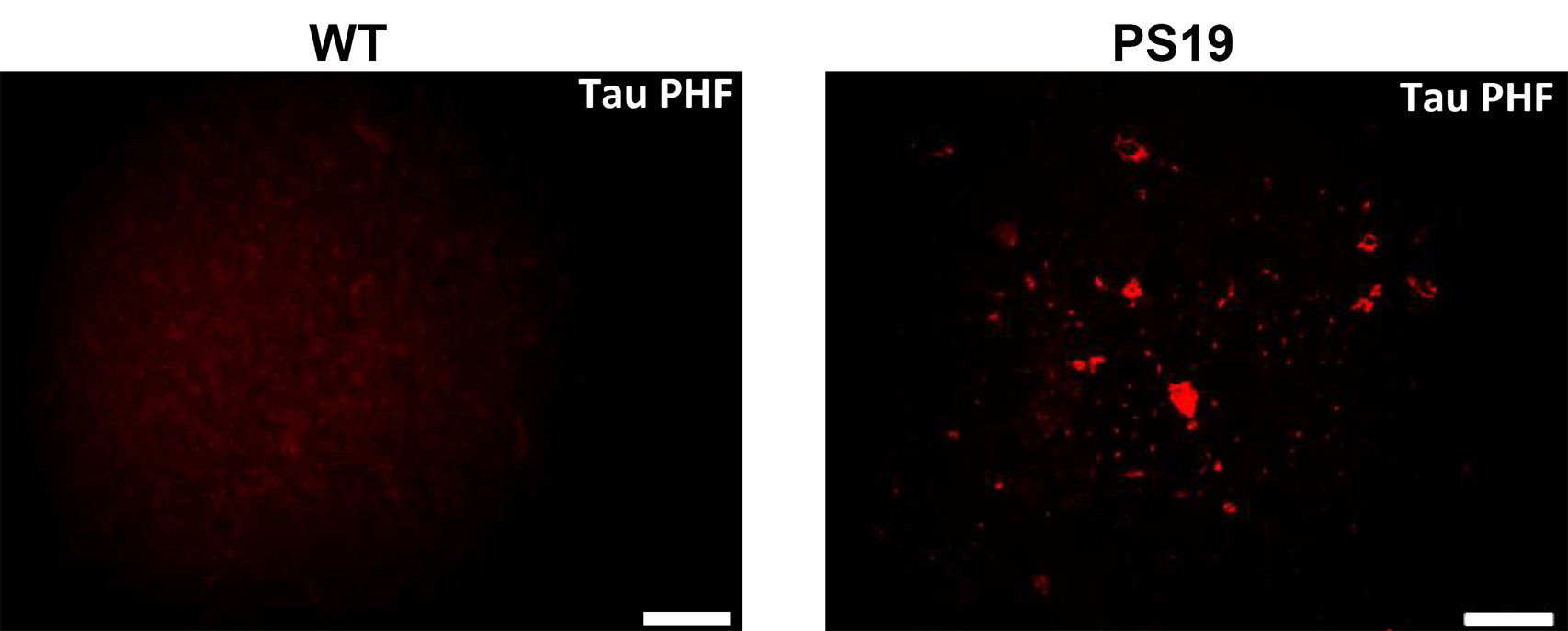
Representative images of PHF1-Tau staining in WT and PS19 brain sections, at 7.5 months. Scale bar=100μm.

**Figure S6.**
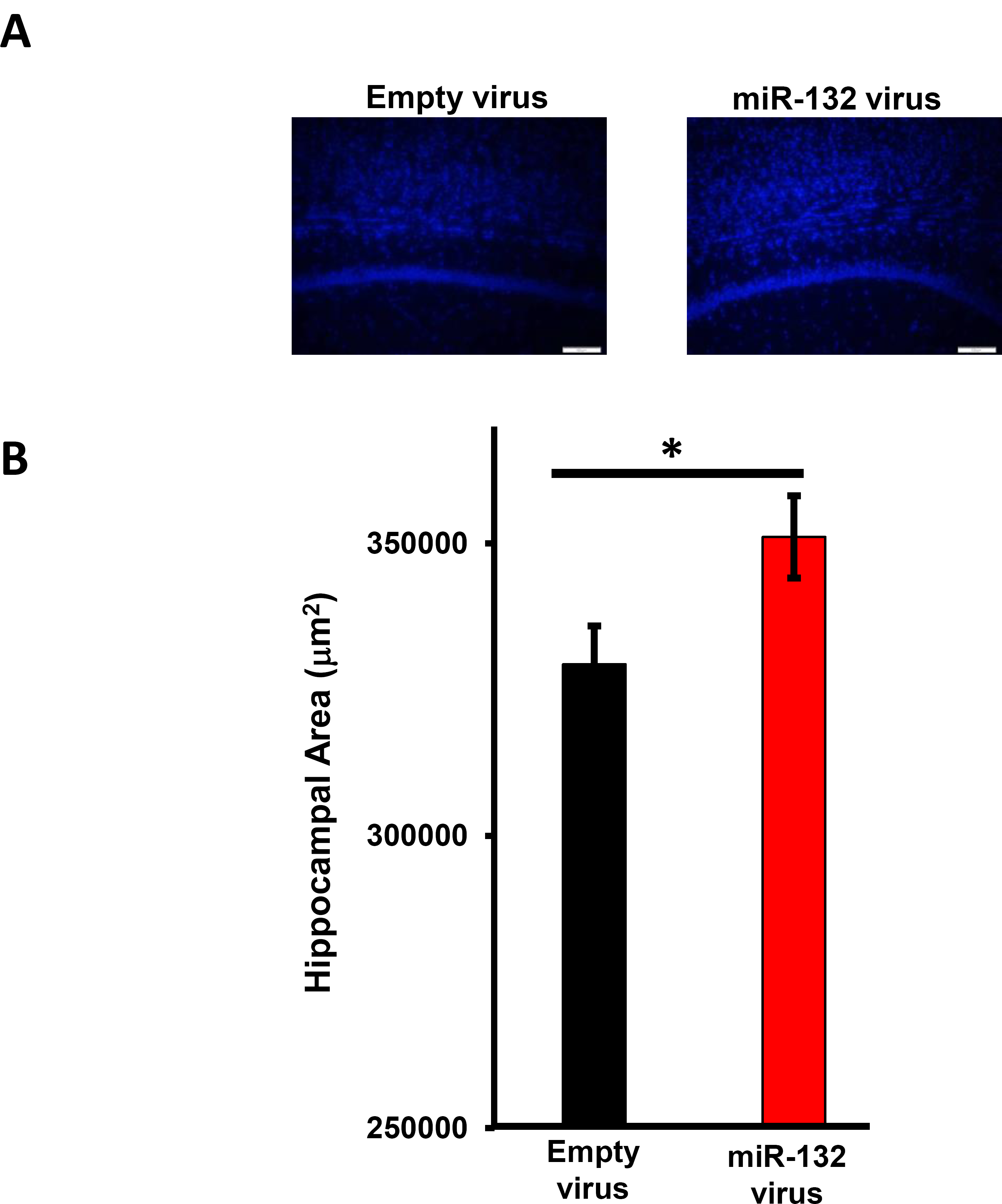
Stereotactic injections of the LV-miR132 to the CA1 reduces hippocampal atrophy in the PS19 mice. **(A)** DAPI staining of PS19 mouse hippocampal CA1 region, injected with the LV-miR132 or control, (**B**) Image J quantification of the hippocampal area (mm^2^), in the brains injected with either LV-miR132 or control virus. For all quantifications, n = 14 mice, 15 sections per brain. All graphical data are shown as mean +/- SEM, Student’s *t*-Tests, *P<0.01.

**Figure S7.**
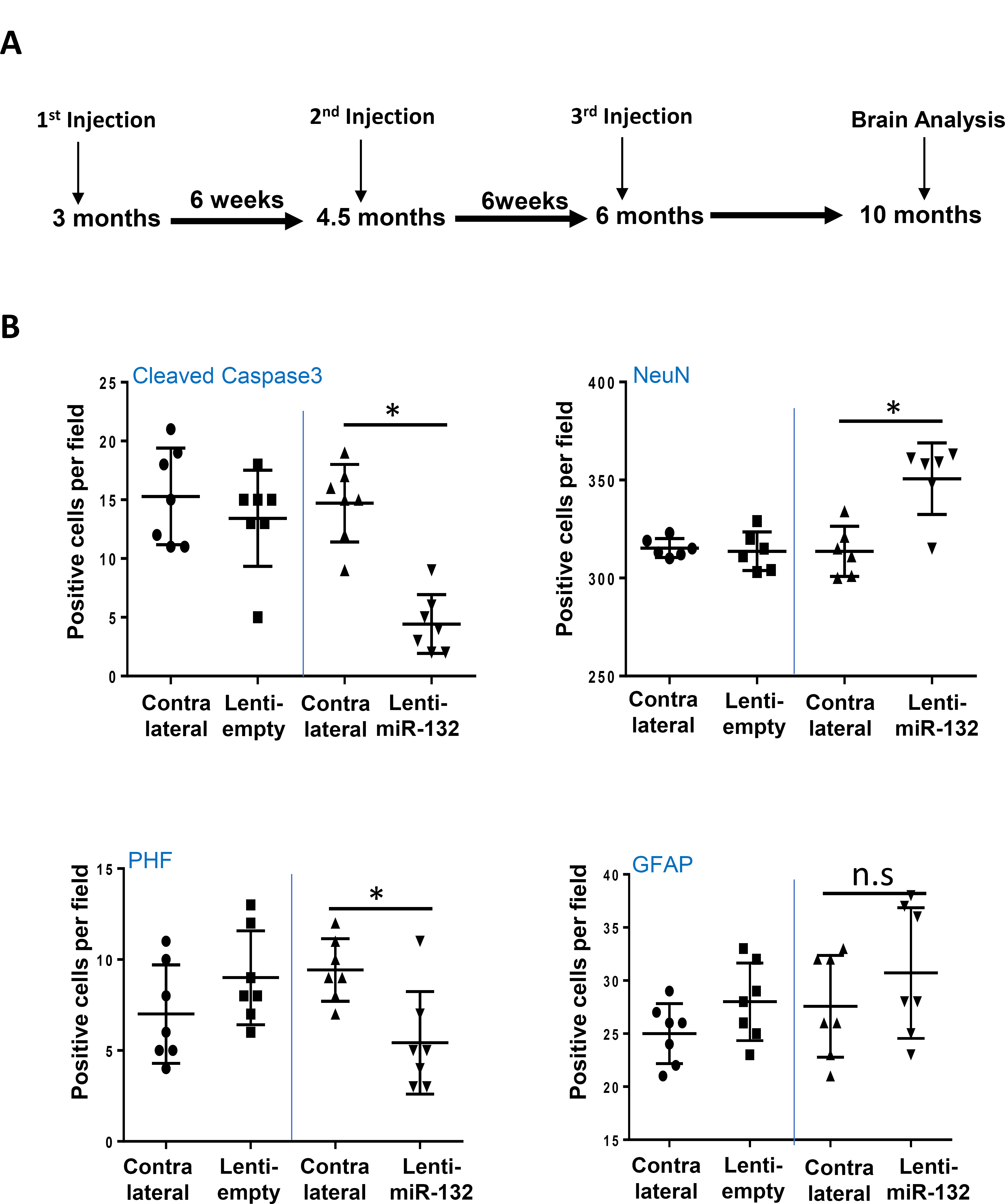
Early supplementation of miR-132 prevents neuronal loss and Tau pathology in PS19 mice. **(A)** Timeline of LV-miR132 injections to young PS19 mice and brain analysis. (**B**) Image J quantification of cells positive for cleaved caspase-3, NeuN, PHF-Tau and GFAP in CA1 and adjacent cortical layers. For all quantifications, n = 7 mice, 15 sections per brain. All graphical data are shown as mean +/- SEM, Student’s *t*-Tests, *P<0.005.

---

**Table S1:**
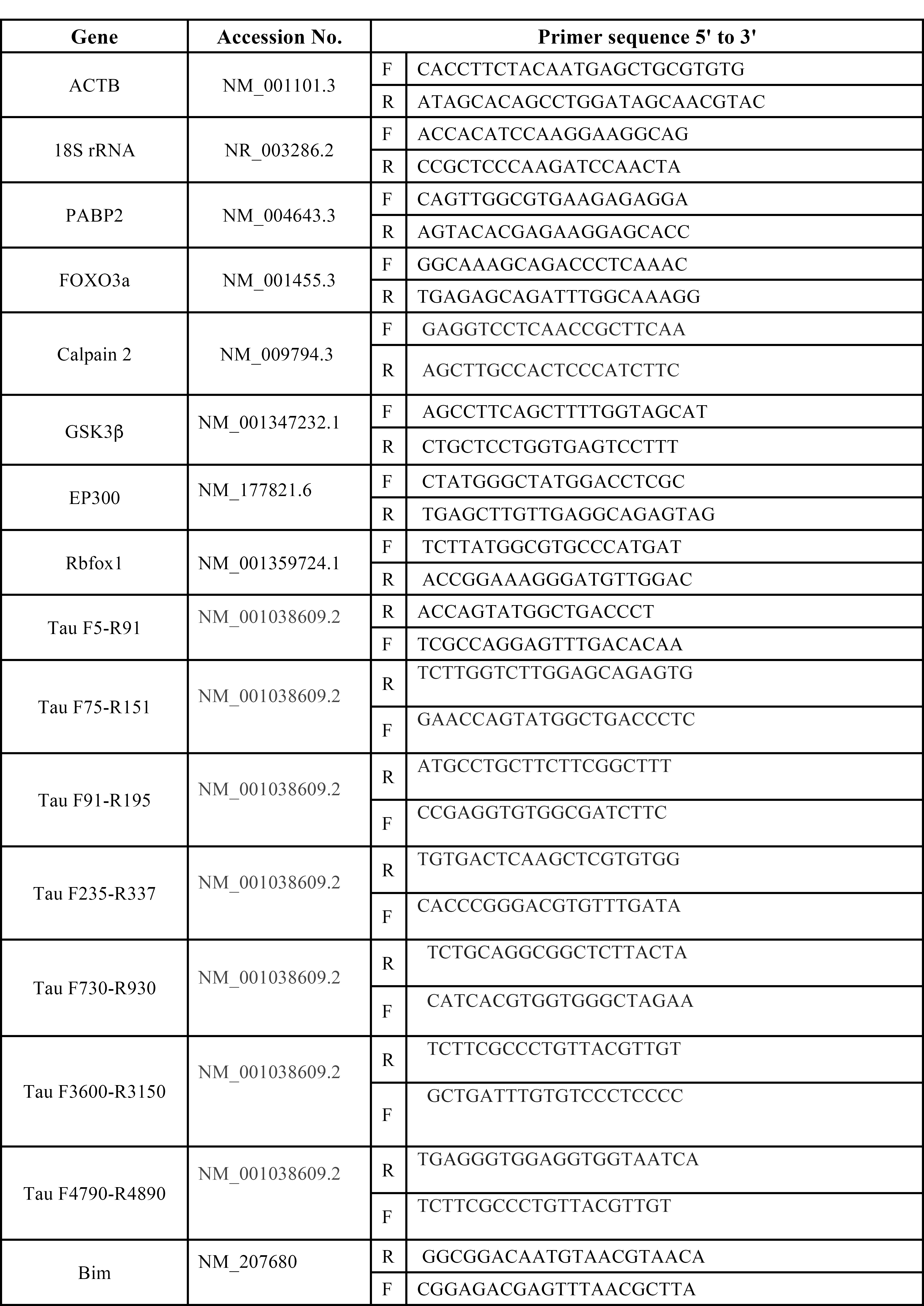
List of primers used for real-time PCR experiments.

